# Natural and pathological aging distinctively impact the vomeronasal detection system and social behavior

**DOI:** 10.1101/2022.08.31.505522

**Authors:** Adrián Portalés, Pablo Chamero, Sandra Jurado

**Author notes:** Correspondence: Dr. Sandra Jurado, Institute of Neuroscience CSIC-UMH, San Juan de Alicante, 03550, Alicante, Spain.

## Abstract

Normal aging and many age-related disorders such as Alzheimer’s disease cause deficits in olfaction, however it is currently unknown how natural and pathological aging impact the detection of social odors which might contribute to the impoverishment of social behavior at old age further worsening overall health. Here, we investigated the effect of aging in the recognition of social cues and the display of social behavior. Our findings indicate that aging distinctively disrupts the processing of social olfactory cues decreasing social odor exploration, discrimination and habituation in both wild type senescent (2-year-old) mice and in 1-year-old double mutant model of Alzheimer’s disease (APP/PS1^Het^). Furthermore, social novelty was diminished in 1-year-old APP/PS1^Het^ mice, indicating that alterations in the processing of social cues are accelerated during pathological aging. Analysis of the vomeronasal organ, the main gateway to pheromone-encoded information, indicated that natural and pathological aging distinctively reduce the neurogenic ability of the vomeronasal sensory epithelium. Cell proliferation remained majorly preserved in 1-year model of Alzheimer’s disease (APP/PS1^Het^), whereas naturally aged animals exhibited significant deficiencies in the number of mature, proliferative and progenitor cells. This study reveals fundamental differences in the cellular processes by which natural and pathological aging disrupt the exploration of social cues and social behavior.

## BACKGROUND

Social recognition is essential for survival allowing to appropriately adapt behavior across a variety of contexts (Adolphs, 2001, Brennan and Kendrick, 2006). As aging progresses, social odors have been shown to elicit attenuated responses (Hlinák and Krejcí, 1991, Garrat et al., 2011), a phenomenon interpreted as a natural consequence of age-related decline in sensory perception (Doty et al., 1984, Murphy et al., 2002, Doty and Kamath, 2014). In humans, social impoverishment has been identified as a major aggravating factor for decreased life expectancy, and an indicator of the appearance of dementia and neurodegenerative disorders such as Alzheimer’s disease (AD) (Heinrichs et al., 2003, Wilson et al., 2007, Donovan et al., 2016). Despite the central role of social interaction in maintaining overall well-being, the mechanisms by which aging, natural or pathological, alter social information processing remain unclear.

Mouse social communication strongly depends on chemosignals comprised by volatile and non-volatile molecules (pheromones) (Beynon and Hurst, 2004, Yoon et al., 2005, Noack et al., 2010, Chamero et al., 2012, Li and Dulac, 2018). Volatile odors are primarily detected at the main olfactory epithelium (MOE) which projects to the main olfactory bulb (MOB), whereas pheromones are mainly detected by the vomeronasal sensory epithelium (VSE) (Jacobson, 1813, Halpern and Martínez-Marcos, 2003, Munger et al., 2009) which connects to the accessory olfactory bulb (AOB) (Ennis et al., 2015). Despite the segregated connectivity, both systems have overlapping functions (Brennan and Zufall, 2006, Zufall and Leinders-Zufall, 2007, Baum and Kelliher, 2009).

A remarkable property of the VSE is that contains proliferative niches capable of generating functional neurons during adulthood (Graziadei and Monti Graziadei, 1983, Weiler et al., 1999, Whitman and Greer, 2009, Brann and Firestein, 2010, Oboti et al., 2015). The neurogenic properties of olfactory structures are known to decline with age, a phenomenon proposed to underlie the olfactory deficits associated to both natural aging and neurodegenerative disorders (Doty et al., 1984, Murphy et al., 2002, Ahlenius et al., 2009, Doty and Kamath, 2014, Mobley et al., 2014, Devanand, 2016). However, how natural and pathological aging affect the regenerative capacity of the VSE has been scarcely studied despite being central to socio-sexual cue detection. An elegant study by Brann and Firestein (2010) showed that the proliferative capacity of the marginal region of the VSE of 2-year-old animals was attenuated in comparison to young animals. More recently, Mechin et al. (2021) reported a significant reduction in mature Olfactory Marker Protein (OMP)-expressing cells (OMP^+^), indicating structural modifications of the aged VNO. However, it is currently unclear whether these cell alterations have any impact in odor detection and social behavior, particularly in animal models of neurodegeneration.

To gain insight into these questions, we investigated the impact of natural and pathological aging in social odor-evoked sniffing behavior, social-odor habituation/dishabituation and social behavior during the aging process of wild type C57/BL6 mice and amyloid precursor protein (APP) and presinilin-1 (PS1) double transgenic mice (APP/PS1^Het^ mice, therein), a widely used animal model of AD (Jankowsky et al., 2001, Jankowsky et al., 2004). Analysis of the number of progenitors, proliferative and mature neurons revealed that naturally aged animals show reduced neuronal proliferation and decreased levels of sensory mature OMP^+^ neurons. In contrast, the VSE of 1-year-old APP/PS1^Het^ mice exhibited normal levels of mature neurons and cell regeneration, suggesting that natural and pathological aging distinctively impact the neurogenic capacity of the VSE.

We explored the functional consequences of these cell alterations observing reduced exploration time of social odors and decreased performance in a social odor habituation-dishabituation test in both senescent and middle age APP/PS1^Het^ mice, suggesting that despite the overall normal VSE structure and cell composition, APP/PS1^Het^ animals exhibit an accelerated decay of social cue decay. Moreover, 1-year-old APP/PS^Het^ mice exhibited a significant reduction of social novelty which was no apparent in naturally aged animals, exposing fundamental differences in how natural and pathological aging impact the regulation of social behavior.

## METHODS

### Animals

Experiments were performed in either control C57/BL6 mice (young: 2-4 months-old; adult: 6-8 months-old; middle age: 12-14 months-old; old (senescent): 20-24 months-old) or APP/PS1 mice on a C57/BL6 genetic background (young: 2-4 months-old; middle-age: 12-14 months-old) (Jackson Labs, Stock No. 004462, MMRRC Stock No. 34829). APP/PS1^Het^ mice express a chimeric mouse/human amyloid precursor protein (Mo/HuAPP695swe) and a mutant human presenilin 1 (PS1-dE9). These animals develop beta-amyloid deposits at ∼ 6 months of age and exhibit early-onset cognitive impairments (Jankowsky et al., 2001, 2004, Reiserer et al., 2007). Animals were kept group-housed and experimentally-naïve during all their lifetime under pathogen-free conditions. Animals were housed in ventilated cages with free access to food and water on a 12 h light/dark cycle.

All experiments were performed according to Spanish and European Union regulations regarding animal research (2010/63/EU), and the experimental procedures were approved by the Bioethical Committee at the Instituto de Neurociencias and the Consejo Superior de Investigaciones Científicas (CSIC).

### VNO dissection

Mice were perfused transcardially with PBS, pH 7.4, followed by 4% paraformaldehyde in 0.1 M phosphate buffer (PB), pH 7.4. A fixed mouse head was placed under a scope and the lower jaw was removed to get a palate view. To facilitate VNO extraction, the palate and the nasal septum were removed and the bilateral VNOs were split in two parts with a gentle twisting motion. Finally, the cartilaginous tissue that covers each portion was carefully removed to extract the VNOs. Tissue was incubated in a 30 % sucrose solution for cryoprotection and kept at −4 °C until sectioning. Samples were embedded in OCT and frozen at −80 °C for cryosectioning.

### Immunohistochemistry

VNOs embedded in OCT-medium were sliced in 16 µm thick sections in a cryostat apparatus (Leica CM 3050S). Slices were incubated in blocking solution containing 0.5 % Triton X-100 and 5 % normal horse serum in 0.1 M TBS for 2 h at room temperature (RT). Primary antibody incubation was performed overnight (o/n) at 4 °C against OMP (Wako goat polyclonal; 1:2000), PCNA (Sigma, rabbit monoclonal; 1:2000); Sox2 (R&D Systems, goat polyclonal; 1:300). For PCNA immunostaining, slices were incubated in 10 mM citrate buffer (100 °C; pH, 6.0) for 5 minutes prior staining. Sections were incubated with Alexa Fluor 488- or 594-conjugated secondary antibodies (Jackson Laboratories, 1:500) for 2 h at RT. Hoechst (Sigma, 1:10000) was added during 5 minutes after the secondary antibody incubation for nuclei visualization. Imaging was performed using a vertical confocal Microscope Leica SPEII. Final images were assembled in Adobe Illustrator.

### Sox2 fluorescence intensity at the VNO supporting cell layer (SCL)

For determining the Sox2 fluorescent labeling area in the VNO SCL where single-cell quantification was limited by the densely packed disposition of the cells, we calculated the corrected total fluorescence (CTF) of the area of interest employing the Freehand ROI tool of Image J. CTF was calculated by subtracting the background fluorescence from a minimum of 3 sections, 16 µm thick, from at least 4 animals per condition.

### Stereology, cell quantification and AOB volume estimation

The total number of olfactory mature neurons (OMP^+^ neurons) in the entire VSE was estimated using stereology (optical fractionator method, Wong et al., 2018) employing Stereo Investigator (MBF Bioscience). Two different tools are combined in this unbiased quantification method: a 3D optical dissector for cell counting and a fractionator, which is based on a systematic, uniform and random sampling over a known area of the tissue (West and Gundersen, 1990, Parrish-Aungst et al., 2007, Dorph-Petersen and Lewis, 2011). The number of neurons was estimated as:

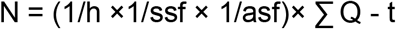

In which **∑ Q** is the total cell number in the region of interest acquired using the optical dissector method; **t** stands for the mean mounted section thickness; **h** is the optical dissector height; **asf** is the area sampling fraction and **ssf** is the frequency of sampling (section sampling fraction). For the VSE stereological analysis, sampling was performed with a 20x objective (Leica, NA 0.6), the counting frame area was 2500 µm^2^ and the sampling grid area was 22500 µm^2^. H for VSE stereology was 8 µm with 1 µm as upper and lower guard zones and t was set at 10 µm. Quantifications were performed for marginal regions of the VSE dividing the area in segments of equal length as previously described (Giacobini et al., 2000, Brann and Firestein, 2010). For the cell quantification in the anterior-posterior regions, the VSE was distributed in 10 sections per animal. Results were divided by the area to obtain the number of cells/mm^2^ in each subdivision. Quantification of the cell number in the marginal area was performed in three slices of each animal delineated with a 50×50 dissector employing Stereo Investigator (MBF Bioscience).

VSE volume was calculated by multiplying the sampled area by the slice thickness (16 µm) and the number of series (10 slices per animal). Ten sections per series were analyzed to obtain an estimation of the VSE area and volume in young (4 months-old), aged (24 months-old), middle aged (12-months old) APP/PS1^WT^ and middle aged APP/PS1^Het^ mice.

The volume of the AOB was estimated by measuring the AOB area and multiplying the sampled area by the section thickness (50 µm) and the number of analyzed sections (5 series per animal).

### Odorants

Social and non-social scents were employed to discern the impact of aging in social odor perception. As non-social scents, we used **i)** the banana-like odor isoamyl acetate (IA) (Sigma) shown to be primarily detected at the MOE level (Xu et al., 2005) with neutral valence within a broad dilution range (Root et al., 2014, Fortés-Marco et al., 2015, Saraiva et al., 2016), and **ii)** food pellets to perform a food finding test (FFT, see experimental details below). As a social scent, we used mouse urine as urine from conspecifics has been shown to elicit robust VNO activity (Chamero et al., 2017). Urine samples were collected according to standard procedures (Kurien et al., 2004). For habituation-dishabituation tests, frozen urine samples from different young subjects and cage mates were used to test fine odor discrimination. For sensitivity tests, urine samples were combined in a stock sample used for each round of odor presentations until completing the experiments. For female urine, samples obtained at different points of the estrous cycle were pooled to generate a uniform stock.

### Behavioral assessment

All the behavioral experiments using social and non-social stimuli were performed in a dedicated room with continuous air reposition under dim indirect light (20 lx). Odor dilutions were prepared in a room outside the animal house. Odor presentation was performed in a homemade methacrylate box with removable walls for cleaning. The chamber dimensions were 15 cm width, 15 cm length and 30 cm high. A small hole (1cm diameter) was performed in the middle of one of the sides located 5 cm above from the box ground to fit standard cotton sticks impregnated with 1 µL of the odorant (**Fig. 3A**) allowing direct contact with the stick. Habituation to the testing conditions was performed before odorant presentations consisting of: handling (5 min), free exploratory activity in the box and habituation to the cotton stick movement (5 min). Odors were presented embedded in a cotton stick in order to preserve non-volatile pheromones and mimic the natural detection method of physical contact between conspecifics (Leypold et al., 2002, Stowers et al., 2002, Kimchi et al., 2007). To avoid variability due to the intensity of the volatile components of the urine, only direct nose contact with the tip of the cotton stick were considered as explorative behavior (sniffing/exploration time). Experiments were monitored by a video recording camera fixed 15 cm above the box. Videos were collected and analyzed off-line using SMART video-tracking software (PanLab S.L.).

#### a) Odor exploration test

After a period of habituation (10 min), mice were exposed to two control trials (mineral oil for isoamyl acetate assays or water for urine tests) during 1 min separated by intervals of 1 min to avoid lack of novelty and odor habituation (Breton-Provencher et al., 2009, Sanderson and Bannerman, 2011). Subsequent presentations consisted on serial dilutions of either urine samples (diluted in water: 1:1.000, 1:500, 1:250, 1:100, 1:50, 1:10 and non-diluted, nd) or isoamyl acetate were tested in ascending order (diluted in mineral oil: 1:5×10^5^, 1:100.000, 1:10.000; 1:1.000, 1:100). Animals were considered to detect the olfactory stimulus when spent more time investigating the odors than the vehicles (water or mineral oil).

#### b) Food finding test (FFT)

Food finding test (FFT) was performed following standard procedures (Deacon et al., 2009). The mice were food deprived for 24 h before testing. Five food pellets (∼ 35 g) were placed a corner and covered by 5 cm of litter bedding. Animals were considered to detect the food pellet when spent digging, touching and holding the food pellet for more than 5 s.

#### c) Social odor habituation-dishabituation test

The effect of aging in social odor discrimination and habituation was explored using an adapted habituation-dishabituation test (Gheusi et al., 2000). After a period of cage habituation (10 min), two control trials were performed employing cotton sticks soaked with 1µl of water (vehicle). Three successive presentations of urine from an animal of the opposite sex (S1a-c) were followed by three presentations of urine from a different subject of the opposite-sex (S2a-c) to evaluate fine chemo-olfactory discrimination and habituation. Urine samples came from young animals of the opposite sex to maximize approaching behavior (Xu et al., 2005, Garratt et al., 2011). Each sample presentation lasted 1 min separated by 1 min intervals. Habituation was estimated as a decrease in the exploration time (sniffing) over the cotton stick tip after the first exposure of urine from the same animal (S1a). Dishabituation (discrimination) occurs in response to a new odor presentation (S2a) as a measurable increase in the exploration time. The test allows to explore two consecutive phases of discrimination (H_2_O^B^-S1a and S1c-S2a) and habituation (S1a-S1b and S2a-S2b). Trend lines between H_2_O^B^-S1a, S1c-S2a, S1a-S1b and S2a-S2b were fitted to obtain the slope values indicating the amplitude of the discrimination and habituation effect in each tested condition. Higher positive values indicated more pronounced social discrimination and higher negative values corresponded to more robust habituation (**Fig. 6**).

#### d) Long-term social habituation-dishabituation test

The aforementioned social odor habituation task was adapted to assay social cue memory by presenting the urine sample employed in S1a, 24 hours after the first test was performed. Intact social odor memory manifested as a reduction in the sniffing time during the presentation of S1a during the second trial, an effect which was clearly apparent in young animals (**Fig. 5F, G**).

#### e) Three chamber test

Social testing was performed in a cage (60 × 40 ×22 cm) following standard procedures (Nadler et al., 2004). Dividing walls were made from clear Plexiglas, with openings allowing access into each chamber. The test mouse was first placed in the middle chamber and allowed to explore for 10 minutes. After the habituation period, an unfamiliar subject of the same sex (mouse 1, M1), was placed in one of the side chambers. The stranger mouse was enclosed in a small, round wire cage, which allowed nose contact between the bars. In this first session (sociability), the test mouse had a choice of spending time in either the empty chamber (E) or the chamber occupied by M1. At the end of the sociability session, each mouse was tested in a second 10-minute session to evaluate social preference for a new subject. A second, unfamiliar mouse (mouse 2, M2) of the same sex was placed in the chamber that had been empty during the first session. This second stranger was enclosed in an identical wire cage than M1. The test mouse had a choice between the first, already-investigated mouse (M1), and the novel unfamiliar mouse (M2) which indicates their social preference or social novelty (Nadler et al., 2004). Continuous video recordings were collected and analyzed off-line using BORIS (Friard and Gamba, 2016) and SMART video-tracking software (PanLab S.L.). Measures of time spent sniffing E, M1, and M2 were quantified for each session.

### Data analysis

All data were tested for statistical significance using GraphPad Prism 8. Shapiro-Wilk test was used to determine data normality. One-way ANOVA with Tukey’s test for multiple comparisons with a single variable was implemented for cell quantifications and anatomical data. For the behavioral analysis, a two-way ANOVA with Tukey’s test was used to test multiple comparisons with more than one variable. P ≤ 0.05 was considered statistically significant. **Additional files 2, 3, 5, 6** summarize the P and N values corresponding to all the statistical comparisons between different ages, genotypes and dilutions of the social odor sensitivity tests.

## RESULTS

### Structural modifications of the mouse VSE during natural and pathological aging

We sought to investigate the structure of the VSE during natural and pathological aging, as the main gateway of pheromone (non-volatile)-encoded information (Sharrow et al., 2002, Cheetham et al., 2007, Ferrero et al., 2013). Stereological analysis revealed significantly smaller VSE volumes in 2 year-old (senescent) mice (**Fig. 1A**: VSE volume mm^3^: 4 month-old: 0.190 ± 0.008; 1-year-old APP/PS1^WT^: 0.170 ± 0.007; 24 month-old: 0.160 ± 0.007; 1-year-old APP/PS1^Het^: 0.160 ± 0.008). This observation was supported by the reduced number of olfactory marker protein positive cells (OMP^+^) along the anterior and medial axis of the marginal VSE (Halpern and Martínez-Marcos, 2003) (**Fig. 1C, E, Additional file 1**: number of OMP^+^ cells ×10^3^ /µm^2^: **Anterior**: 4 month-old: 10.69 ± 0.56; 12 month-old APP/PS1^WT^: 9.50 ± 0.45; 24 month-old: 7.80 ± 0.53; 12 month-old APP/PS1^Het^: 8.60 ± 0.53; **Medial**: 4 month-old: 9.92 ± 0.23; 12 month-old APP/PS1^WT^: 9.15 ± 0.78; 24 month-old: 7.70 ± 0.70; 12 month-old APP/PS1^Het^: 8.40 ± 0.40; **Posterior**: 4 month-old: 9.15 ± 0.70; 12 month-old APP/PS1^WT^: 9.53 ± 0.77; 24 month-old: 7.90 ± 0.75; 12 month-old APP/PS1^Het^: 10.00 ± 0.84).

**Figure 1.**
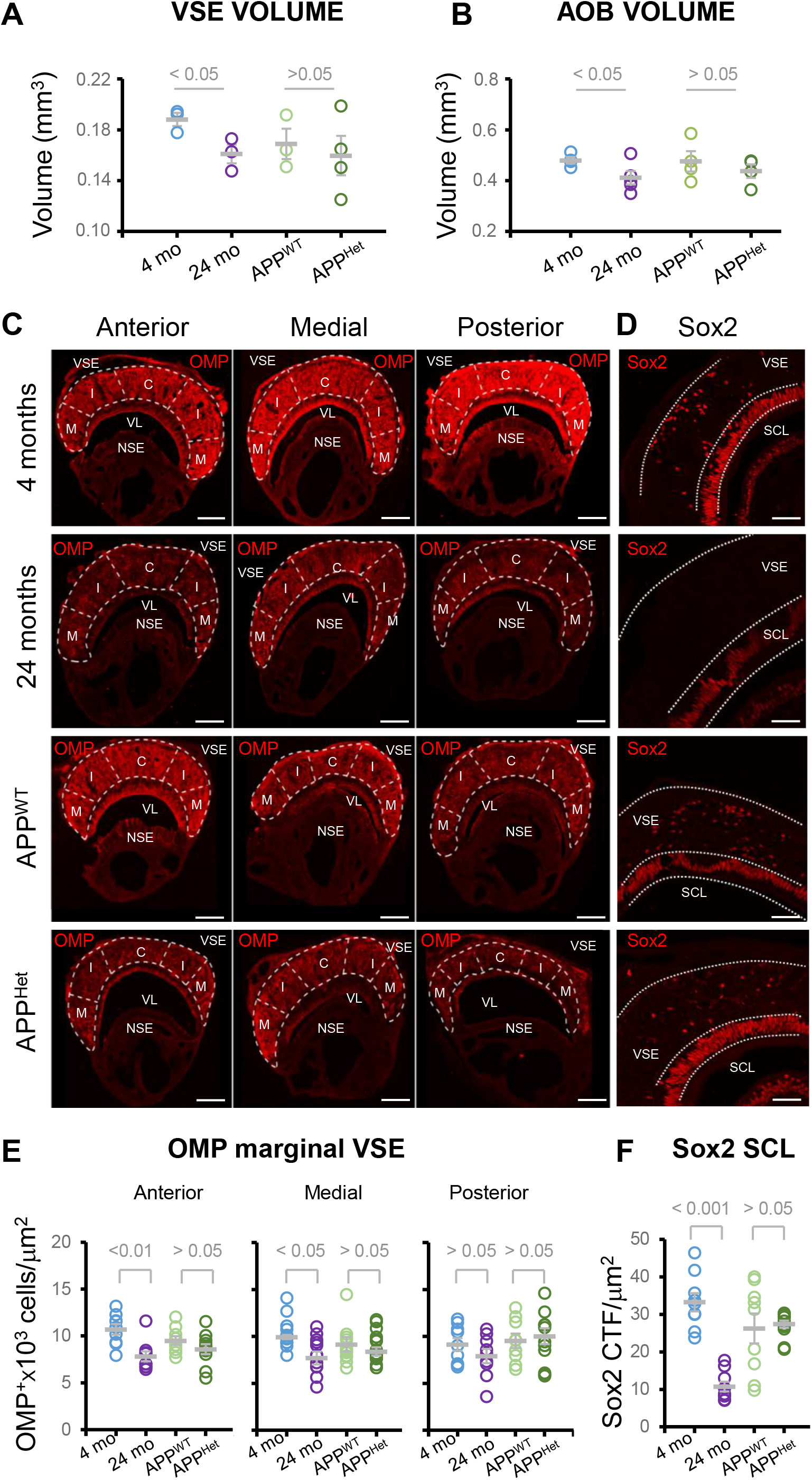
Structural modifications of the mouse VSE during natural and pathological aging. **A)** Dispersion plot of the VSE volume indicates a significant reduction during natural but not pathological aging. **B)** Dispersion plot of AOB volume. Thick lines indicate the mean ± SEM. Data were obtained from 4 animals per condition. **C)** Representative images of OMP staining in the VSE during natural and pathological aging. Scale bar indicates 100 µm. **D)** Representative images of Sox2 staining in the VSE and the supporting cell layer. Scale bar indicates 100 µm. VL: vomeronasal lumen; SCL: supporting cell layer; NSE: non-sensory epithelium; M: marginal zone; I: intermediate zone; C: central zone. **E, F)** Dispersion plots represent the number of OMP^+^ cells in the VSE (number of cells × 10^3^/µm^3^) and the intensity of Sox2 fluorescence in the SCL (Sox2 CTF /µm^2^). Thick lines indicate the mean ± SEM. Data were obtained from 10-16 slices from at least three animals per condition. Statistical comparisons were calculated by two-way ANOVA with Tukey’s test. P ≤ 0.05 was considered statistically significant.

We hypothesized that the drop in the number of OMP^+^ cells could translate into a reduced axonal projection to the AOB, the main VNO target area, resulting in smaller AOB volumes. As such, we observed reduced AOB size in 2 year-old mice (**Fig. 1B:** AOB volume mm^3^: 4 month-old: 0.48 ± 0.03; 12 month-old APP/PS1^WT^: 0.48 ± 0.04; 24 month-old: 0.41 ± 0.03; 12 month-old APP/PS1^Het^: 0.44 ± 0.03), indicating that aging may alter basic structural features of the VNO-AOB axis.

We expanded our analysis to the SRY-box transcription factor 2 (Sox2)-expressing cells (Sox2^+^) in the supporting cell layer (SCL), a neuronal stem cell-marker which also stain mature differentiated sustentacular cells (Guo et al., 2010, Taroc et al., 2020, Katreddi et al., 2021). The estimation of the labeling intensity of Sox2 in the SCL showed a drastic reduction in senescent but not in APP/PS1^Het^ mice (**Fig. 1D, F**: Sox2 fluorescence intensity / area (µm^2^): 4 month-old: 33.24 ± 2.43; 12 month-old: 26.23 ± 3.95; 24 month-old: 10.74 ± 1.32; 12 month-old APP/PS1^Het^: 27.37 ± 0.96), indicating natural aging may disrupt VSE structure by reducing the number of sensory neurons and sustentacular cells (OMP^+^ and Sox2^+^ cells, respectively).

### Natural and pathological aging differentially impact VSE cell proliferation

The observed decrease in the number of mature OMP^+^ and Sox2^+^ cells in the VSE of senescent mice could be due to the reduction of the VSE’s regenerative ability. Given that there is currently scarce data on the properties of the VSE neurogenic niche in neurodegenerative animal models, we analyzed the number of proliferative cells identified by the expression of proliferative cell nuclear antigen (PCNA) in the anterior, medial and posterior VNO in senescent and APP/PS1^Het^ mice. Consistent with previous reports cell proliferation in young animals was abundant in the marginal zone of the anterior and medial VSE (Giacobini et al., 2000, Brann and Firestein 2010, **Fig. 2A, B**). According to previous reports (Brann and Firestein, 2010), senescent mice exhibited a significant reduction in PCNA^+^ cells in the marginal VSE (**Fig. 2A, B**, number of PCNA^+^ cells × 10^3^/mm^3^: **Anterior**: 4 month-old: 2.26 ± 0.80; 12 month-old APP/PS1^WT^: 0.73 ± 0.3; 24 month-old: 0.75 ± 0.35; 12 month-old APP/PS1^Het^: 3.42 ± 0.82; **Medial**: 4 month-old: 2.15 ± 0.52; 12 month-old APP/PS1^WT^: 1.61 ± 0.32; 24 month-old: 0.23 ± 0.13; 12 month-old APP/PS1^Het^: 1.57 ± 0.20; **Posterior**: 4 month-old: 3.04 ± 0.90; 12 month-old APP^WT^: 2.42 ± 0.60; 24 month-old: 0.17± 0.08; 12 month-old APP/PS1^Het^: 1.93 ± 0.57), indicating reduced proliferation. Surprisingly, the number of PCNA^+^ cells in APP/PS1^Het^ mice was significantly higher that of aged-matched controls in the anterior VSE (**Fig. 2A, B**), suggesting a region-specific increase in cell generation in these animals. Importantly, no significant overlap between OMP and PCNA staining was observed indicating that these two markers recognize cell populations at different maturation stages (**Fig. 2A**, right panel).

**Figure 2.**
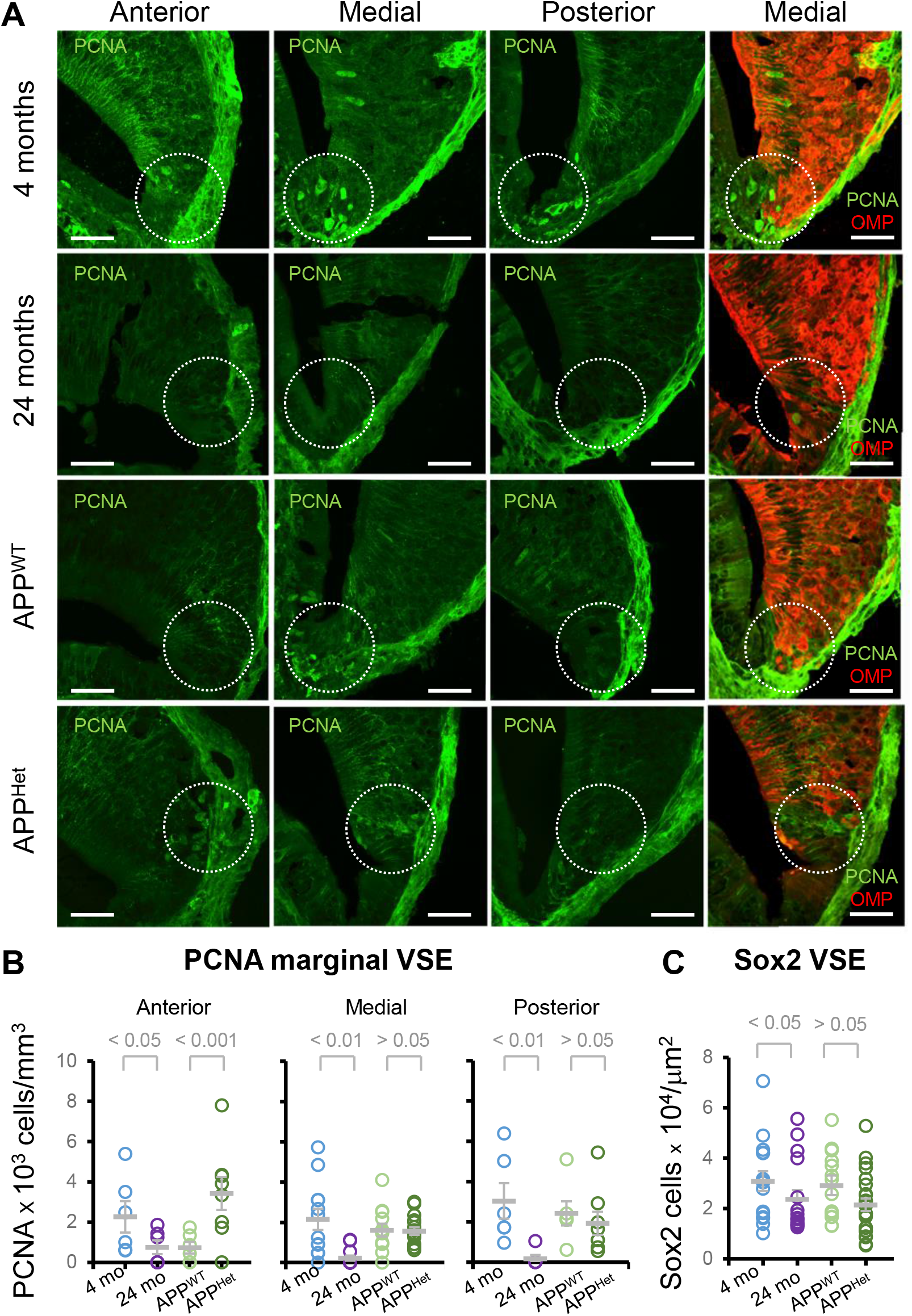
Natural and pathological aging differentially impact cell proliferation in the VSE. **A)** Representative confocal images of PCNA staining in the VSE proliferative niche during natural and pathological aging. Panels on the right show confocal images of OMP and PCNA staining in the VNO. To note there is no overlapping signal between the two markers indicating the identification of cells at different maturation stages. Circles indicate the proliferative marginal region of the VNO. Scale bar represents 20 µm. **B, C)** Dispersion plots represent the number of PCNA^+^ cells (number of cells × 10^3^/mm^3^) and the number of Sox2^+^ cells (number of cell × 10^4^/µm^2^) in the marginal VNO. Thick lines indicate the mean ± SEM. Data were obtained from ten slices from animal of each condition. Statistical analysis was calculated by two-way ANOVA with Tukey’s test for multiple comparisons. P ≤ 0.05 was considered statistically significant.

We then sought to explore whether the reduced number of proliferative PCNA^+^ cells in senescent mice could be due to a decrease of stem cell generation by analyzing the number of Sox2^+^ neural precursor cells in the VSE (Tucker et al., 2010, Panaliappan et al., 2018). We observed a significant reduction in the number of Sox2^+^ cells in 2 year-old mice but not in APP/PS1^Het^ animals, indicating a reduced ability to progenitor generation in the aged VSE (**Fig. 2C**: number of Sox2^+^ cells in the VSE, number of cells × 10^4^ cells/µm^2^: 4 month-old: 3.08 ± 0.40; 12 month-old APP/PS1^WT^: 2.92 ± 0.38; 24 month-old: 2.37 ± 0.36; 12 month-old APP/PS1^Het^: 2.15 ± 0.24). Altogether, these findings revealed a fundamental difference in how natural and diseased aging impact the VSE structure and proliferative capacity.

### Late onset of social exploration deficits during natural aging

Our data indicated that natural aging induce significant alterations in the structure, VSE proliferative capacity and cellular composition, thus we investigated whether these adaptations might translate into impairments in the processing of socio-sexual information. Cumulative evidence points to olfactory decline as a common symptom of natural and pathological aging (Doty et al., 1984, Murphy et al., 2002, Rawson et al., 2012, Doty and Kamath, 2014, Roberts et al., 2016). However, most of these studies and diagnostic tests employed synthetic odors with reduced emotional valence, therefore, the temporal course and severity of the age-related involution of the recognition of social cues, largely processed by the VNO-AOB system, remains unclear.

To explore the impact of natural aging on social odor detection, we conducted an odor-evoked sniffing test in which serial dilutions of urine from young conspecifics of the opposite sex were presented to either male or female subjects over a range of different ages: young (2-4 mo.), adult (6-8 mo.), middle-aged (12-14 mo.) and old/senescent mice (20-24 mo.) (**Fig. 3A**, see Material and Methods for details on urine sample preparation). Our results indicated a significant reduction in the exploration time throughout the various urine dilutions in comparison to adult wild type mice (**Fig. 3B, C** sniffing time normalized to water control 2, **undiluted sample**: aged: 3.83 ± 0.71; adult: 6.06 ± 0.88; **1:10 dilution**: aged: 3.04 ± 0.50; adult: 4.90 ± 1.10; **1:50 dilution**: aged: 1.90 ± 0.33; adult: 3.14 ± 0.55; **1:100 dilution**: aged: 1.56 ± 0.37; adult: 2.95 ± 0.43; **1:250 dilution**: aged: 1.37 ± 0.36; adult: 2.36 ± 0.70; **1:500 dilution**: aged: 0.86 ± 0.20; adult: 2.50 ± 0.35; **1:1000 dilution**: aged: 0.60 ± 0.20; adult: 2.80 ± 0.43). Raw sniffing time data showed a similar decrease in the exploration of social odors in senescent animals across different dilutions (see details in **Additional files 2, 3**).

Furthermore, senescent mice progressively increased the exploration time in parallel with odor concentration, whereas young, adult and middle-age animals showed a habituation phase at intermediate dilutions (1:500, 1:250, 1:100 and 1:50), indicating effective odor detection and recognition capabilities (Yang and Crawley, 2009). Lastly, data analysis disaggregated by sex showed no sex-differences in the reduction of urine exploration time (**Additional file 4**; but see Martínez et al., 2017 for an alternative view).

### Reduction in the exploration of social information is accelerated in an animal model of neurodegeneration

Next, we investigated whether pathological aging would alter the exploration of social olfactory cues despite no obvious effects on VSE structure or proliferative capacity. Social odor sensitivity tests in 1 year-old APP/PS1^Het^ mice revealed reduced sniffing times when compared to age-matched APP/PS1^WT^ control mice for low urine dilutions, suggesting that pathological aging accelerates the decline of exploration time of social odors (**Fig. 3D, E** sniffing time normalized to water control 2, **undiluted sample**: 1 year-old APP/PS1^Het^: 2.58 ± 0.43; 1 year-old APP/PS1^WT^: 5.65 ± 1.51; **1:10 dilution**: 1 year-old APP/PS1^Het^: 1.82 ± 0.27; 1 year-old APP/PS1^WT^: 4.94 ± 1.21; **1:50 dilution**: 1 year-old APP/PS1^Het^: 1.43 ± 0.46; 1 year-old APP/PS1^WT^: 1.40 ± 0.26; **1:100 dilution**: 1 year-old APP/PS1^Het^: 1.37 ± 0.43; 1 year-old APP/PS1^WT^: 2.03 ± 0.55; **1:250 dilution**: 1 year-old APP/PS1^Het^: 0.90 ± 0.17; 1 year-old APP/PS1^WT^: 0.70 ± 0.14; **1:500 dilution**: 1 year-old APP/PS1^Het^: 0.52 ± 0.10; 1 year-old APP/PS1^WT^: 0.97 ± 0.20; **1:1000 dilution**: 1 year-old APP/PS1^Het^: 1.00 ± 0.20; 1 year-old APP/PS1^WT^: 0.84 ± 0.31). Similar to senescent mice, these results were reproduced when raw sniffing time data were compared (**Additional file 5, 6**).

### Aging and neurodegeneration impact social odor recognition more severely than other odor modalities

Next, we asked whether non-social odor modalities were also affected in senescent and APP/PS1^Het^ animals. We investigated this question by analyzing the exploration time to both food and synthetic neutral odors. First, we exposed naturally aged and APP/PS1^Het^ mice to serial dilutions of IA, a synthetic banana like odor of neutral valence when used at high dilutions (Fortes-Marco, 2015, Saraiva et al., 2016). Consistent with previous studies, mice of all conditions showed significantly reduced responses to the neutral odorant than to urine (**Fig. 4A-D**) according to the higher emotional valence of urine in comparison to a synthetic odor (Kobayakawa et al. 2007, Mandairon et al. 2009, Saraiva et al. 2016, Jagetia et al. 2018). Our results showed no significant differences on the exploration times of IA in senescent mice and a modest but significant increase in the sniffing time of the 1:10^4^ and 1:100 IA dilutions in APP/PS1^Het^ animals (**Figures 4A-D**, normalized sniffing time, **1:100 dilution**: 1 year-old APP/PS1^Het^: 1.40 ± 0.12; 1 year-old APP/PS1^WT^: 0.93 ± 0.16; young: 1.76 ± 0.26; aged: 1.24 ± 0.20; 1:1000 dilution: 1 year-old APP/PS1^Het^: 1.40 ± 0.22; 1 year-old APP/PS1^WT^: 1.42 ± 0.30; young: 1.90 ± 0.35; aged: 1.56 ± 0.50; **1:10**^**4**^ **dilution**: 1 year-old APP/PS1^Het^: 1.50 ± 0.25; 1 year-old APP/PS1^WT^: 0.91 ± 0.13; young: 1.44 ± 0.32; aged: 1.20 ± 0.26; **1:10**^**5**^ **dilution**: 1 year-old APP/PS1^Het^: 0.75 ± 0.11; 1 year-old APP/PS1^WT^: 0.75 ± 0.15; young: 1.41 ± 0.50; aged: 0.90 ± 0.20; **1:5×10**^**5**^ **dilution**: 1 year-old APP/PS1^Het^: 0.85 ± 0.18; 1 year-old APP/PS1^WT^: 0.82 ± 0.11; young: 1.65 ± 0.42; aged: 1.23 ± 0.35). Similarly, the FFT showed no significant differences in the latency to find food pellets after 24 hours of food deprivation in either naturally aged or middle age APP/PS1^Het^ mice (**Fig. 4E** latency of finding food pellet (min), young: 5.22 ± 1.08; aged: 5.15 ± 0.95; 1-year-old APP/PS1^Het^: 4.27 ± 0.58; 1-year-old APP/PS1^WT^: 3.87 ± 0.32). These results suggest that deficits in social exploration time at advanced stages of natural aging and in an animal model of AD are more pronounced than other odor modalities.

**Figure 3.**
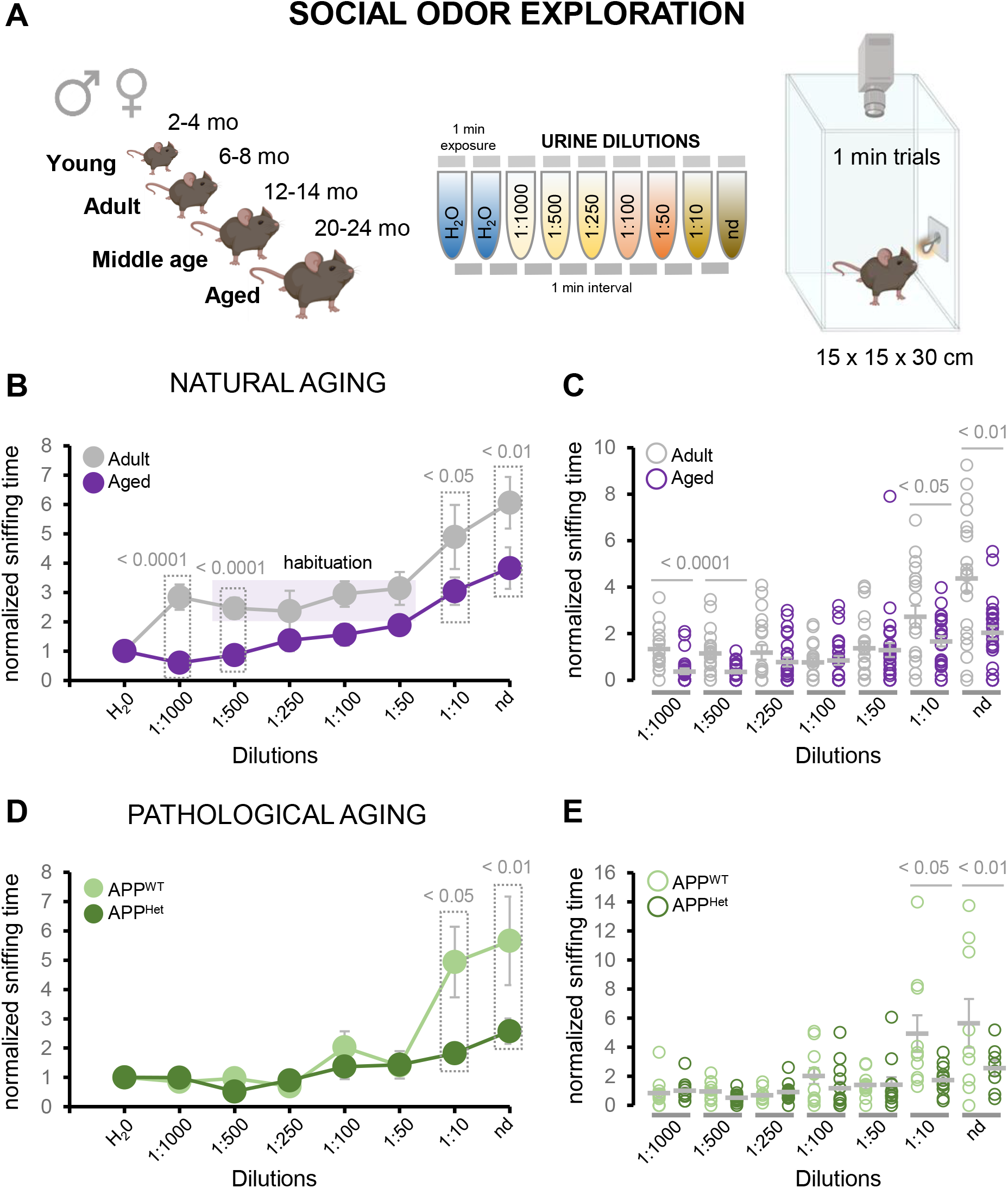
Natural and pathological aging reduce exploration to social odors. **A)** Schematics of the olfactory test used in this study in which urine dilutions are presented as a social signal. **B)** Average of the exploration (sniffing) time of urine serial dilutions normalized to the exploration time of the vehicle (water) of adult and aged wild type mice. A typical habituation was observed in adult mice at intermediate dilutions (1:500, 1:250, 1:100; 1:50). **C)** Dispersion plot of the normalized sniffing time of each urine dilution (nd, non-diluted; 1:10; 1:50; 1:100; 1:250; 1:500; 1:1000) for adult and aged wild type mice. **D)** Average of the sniffing time of urine dilutions normalized to the exploration time of the vehicle (water) of middle age APP/PS1 controls (APP^WT^) and middle age APP/PS1 heterozygous mice (APP/PS1^Het^). **E)** Dispersion plots of normalized sniffing time of middle age APP^WT^ control and APP/PS1^Het^ mice. Grey lines in the dispersion plots indicate mean ± SEM. Data were analyzed by a one-way ANOVA with Tukey’s test to test multiple comparisons with more than one variable. P ≤ 0.05 was considered statistically significant.

**Figure 4.**
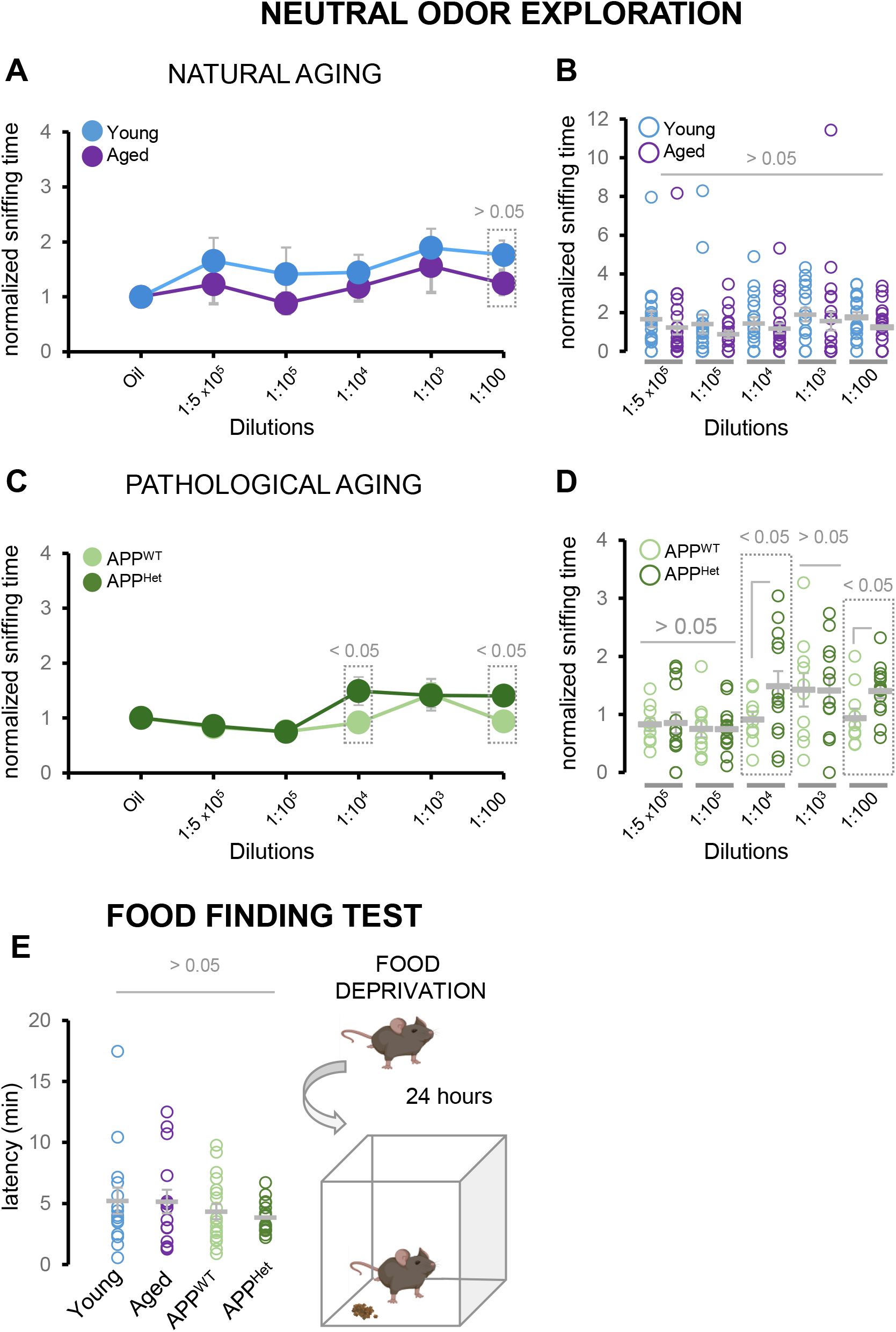
Aging and neurodegeneration impact social odor recognition more severely than other odor modalities. **A)** Average of the sniffing time of young and aged mice in response to a neutral synthetic odor isoamyl acetate (IA) normalized to the vehicle (mineral oil). **B)** Dispersion plots of the normalized sniffing time of young and aged mice in response to various dilutions of IA. **C)** Average of the sniffing time of middle age APP/PS1 controls (APP^WT^) and middle age APP/PS1 heterozygous mice (APP/PS1^Het^) in response to IA normalized to the vehicle (mineral oil). **D)** Dispersion plots of normalized sniffing time of APP/PS1^WT^ and APP/PS1^Het^ mice in response to various dilutions of IA. Note that APP/PS1^Het^ mice exhibit significantly higher exploration times for the 1:10^4^ and 1:100 IA dilutions. **E)** Food deprived animals performed a food seeking test (schematics on the right) which revealed equivalent latencies to the find hidden food pellets as shown in the dispersion data plot (left panel). Thick lines in dispersion plots indicate mean ± SEM. Data were analyzed by a one-way ANOVA with Tukey’s test to test multiple comparisons with more than one variable. Number of mice: *Neutral odor sensitivity*: young: n = 19; aged: n = 27; middle age control: n = 10; middle age APP/PS1^Het^: n = 13. *Food finding test*: young: n = 21; aged: n = 27; APP^WT^: n = 20; APP^Het^: n = 16. P ≤ 0.05 was considered statistically significant.

### Social odor discrimination and habituation are reduced in naturally aged and APP/PS1^Het^ mice

To further investigate the impact of aging and neurodegeneration in the detection of social information, we performed a habituation – dishabituation test, which relies on the animal’s ability to discriminate novel smells (Yang and Crawley, 2009). For these experiments, young (2-4 mo.) and aged (20-24 mo.) wild type animals were presented three consecutive replicates of urine samples from two different animals (S1a-c and S2a-c) (**Fig. 5A**). Aged animals were able to discriminate between urine sources, but showed reduced sniffing times during both the first and second discrimination and habituation phases (**Fig. 5B, C**). Similarly, middle age APP/PS1^Het^ mice also exhibited reduced sniffing times in comparison to aged-matched APP/PS1^WT^ controls (**Fig. 5D, E** sniffing time in seconds, **water control 1**: aged: 0.56 ± 0.09; young: 0.42 ± 0.05; APP/PS1^WT^: 0.72 ± 0.08; APP/PS1^Het^: 0.57 ± 0.06; **water control 2**: aged: 0.56 ± 0.11; young: 0.67 ± 0.10; APP/PS1^WT^: 0.50 ± 0.07; APP/PS1^Het^: 0.71 ± 0.09; **S1a**: aged: 1.36 ± 0.27; young: 2.45 ± 0.40; APP/PS1^WT^: 2.81 ± 0.40; APP/PS1^Het^: 1.75 ± 0.22; **S1b**: aged: 0.70 ± 0.14; young: 0.87 ± 0.20; APP/PS1^WT^: 0.95 ± 0.10; APP/PS1^Het^: 0.68 ± 0.10; **S1c**: aged:0.70 ± 0.21; young: 0.51 ± 0.10; APP/PS1^WT^: 0.84 ± 0.22; APP/PS1^Het^: 0.71 ± 0.13; **S2a**: aged:1.51 ± 0.27; young: 2.44 ± 0.43; APP/PS1^WT^: 1.85 ± 0.42; APP/PS1^Het^: 1.00 ± 0.17; **S2b**: aged: 0.82 ± 0.14; young: 0.81 ± 0.15; APP/PS1^WT^: 0.80 ± 0.22; APP/PS1^Het^: 0.62 ± 0.08; **S2c**: aged: 0.96 ± 0.18; young: 0.55 ± 0.11); APP/PS1^WT^: 0.60 ± 0.13; APP/PS1^Het^: 0.50 ± 0.09). Analysis of the slope values of the first and second discrimination and habituation phases revealed that senescent mice exhibited significant deficits during both rounds of social habituation – dishabituation, and APP/PS1^Het^ mice only during the second phase of discrimination (**Fig. 6A, B**).

Then, we adapted the habituation-dishabituation test to assay potential changes in long-term social odor memory (**Fig. 5F**). To this aim, animals were exposed to the same S1a urine sample 24 hours after the first presentation. A reduction of the sniffing time during the first discrimination phase was interpreted as an indicator of memory. This reduction was clearly observed in young animals, but absent in naturally aged and 1-year-old APP/PS1^Het^ mice (**Fig. 5G:** sniffing time of S1a after 24 hours in seconds, young: 1.20 ± 0.20; aged: 1.12 ± 0.20; APP/PS1^WT^: 1.40 ± 0.50; APP/PS1^Het^: 1.20 ± 0.33), suggesting long-term social odor memory impairments during both natural and pathological aging.

### Age-related deficits in social discrimination and habituation are not influenced by previous experience

Next, we compared animals exposed to either familiar (littermate urine, L) or novel social odors (novel urine, N) in the habituation-dishabituation test (**Fig. 7A**). Aged animals performed poorly in the L-N dishabituation task (no significant increase in sniffing time), suggesting that the reduction in social odor discrimination and habituation is independent of previous experience.

**Figure 5.**
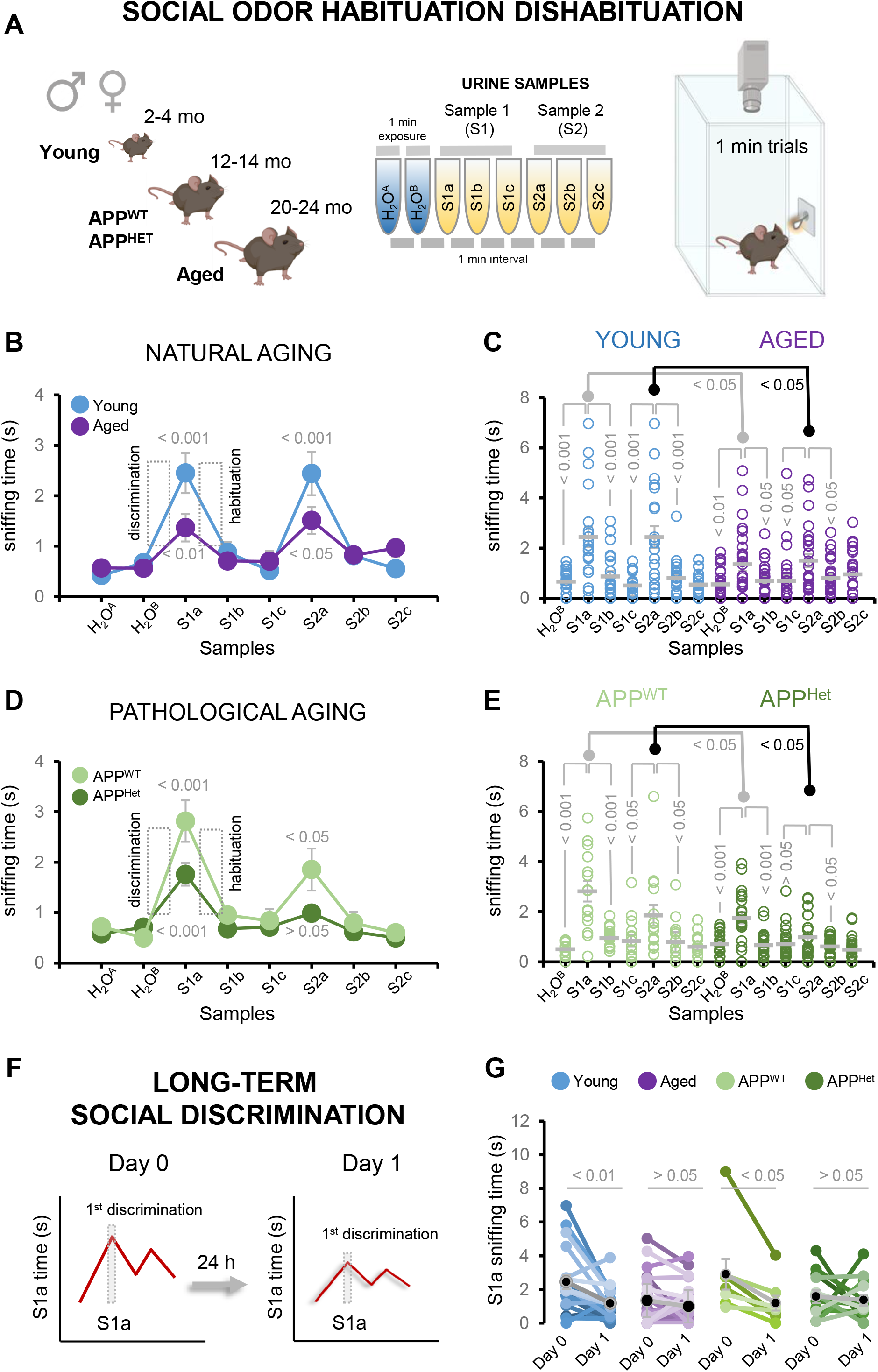
Social odor discrimination and habituation are reduced in naturally aged and APP/PS1^Het^ mice. **A)** Schematics of the social habituation-dishabituation test used in this study: after habituation to the experimental cage, the animal is exposed to the same urine sample three times (S1a-c) which induces a typical increase in exploration time (1^st^ discrimination) to be followed by a reduction in the sniffing time (1^st^ habituation). Dishabituation induced by a urine sample from a new subject (S2a) elicits a second round of discrimination - habituation. **B)** Average sniffing time of the social habituation-dishabituation test performed by young and aged wild type mice. **C)** Dispersion plot of the sniffing time of young and aged animals during the social habituation-dishabituation test. **D)** Average sniffing time of the social habituation-dishabituation test performed by middle age APP/PS1^WT^ control and APP/PS1^Het^ mice. **E)** Dispersion plot of the sniffing time of middle age APP/PS1^WT^ control and APP/PS1^Het^ mice during the social habituation-dishabituation test. **F**) The habituation-dishabituation test was modified to assess potential deficits in long-term discrimination due to natural or pathological aging by presenting the same S1a sample 24 hours after. **G**) Paired data corresponding to the sniffing time during the first and second S1a presentation (24 hours later) is plotted for each condition. Thick lines in dispersion plots and black dots in G indicate mean ± SEM. Data were analyzed by a one-way ANOVA with Tukey’s test to test multiple comparisons with more than one variable. P ≤ 0.05 was considered statistically significant.

**Figure 6.**
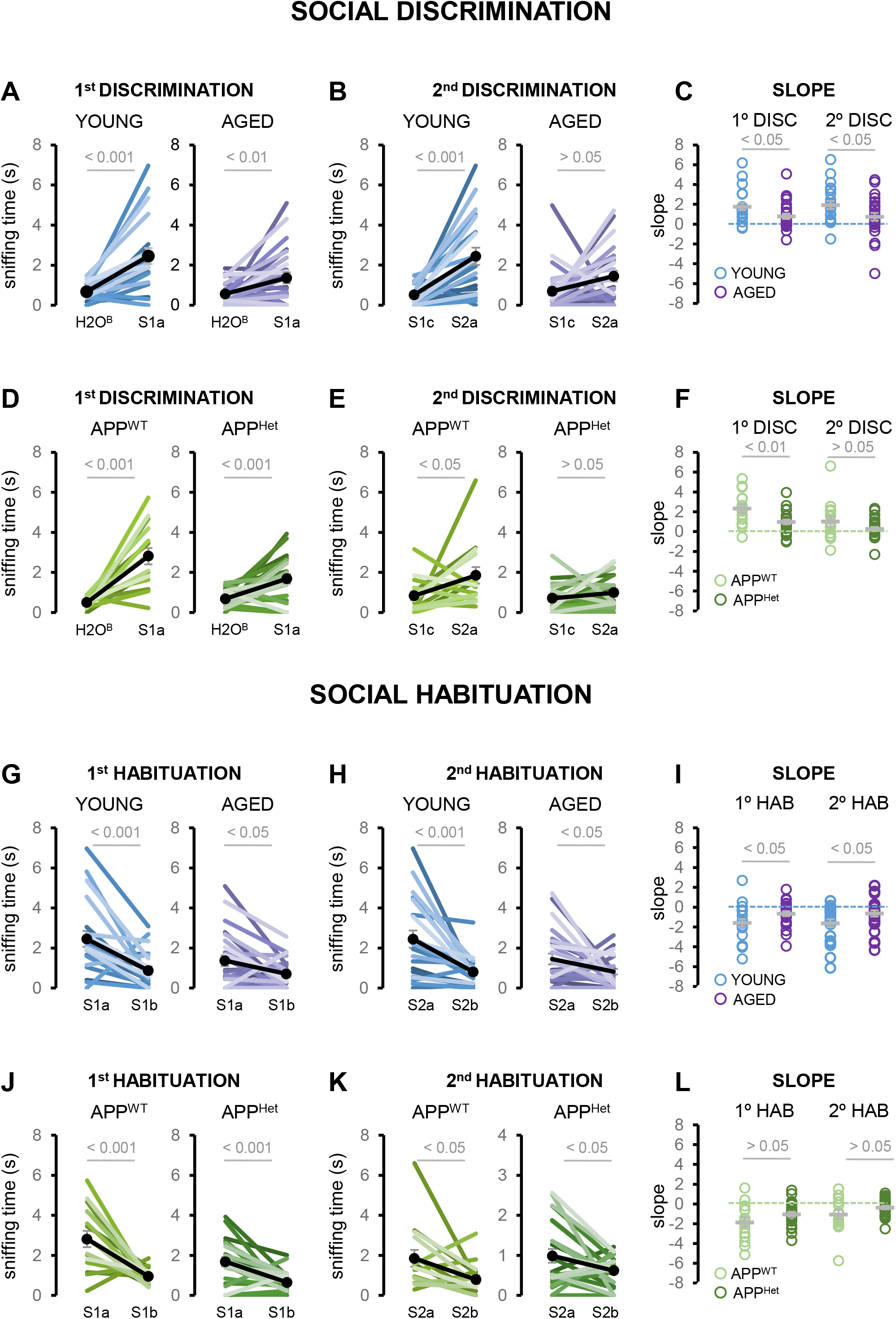
Social discrimination and habituation are more severely altered during natural aging. **A)** Paired data of the sniffing time young and aged wild type animals corresponding to the first discrimination phase (H_2_O-S1a). **B**) Paired data of the sniffing time of young and aged wild type animals corresponding to the second discrimination phase (S1c-S2a). **C**) Dispersion plots of the slope values corresponding to the first and second discrimination phases of young and aged mice. **D)** Paired data of the sniffing time of APP^WT^ and APP^Het^ mice corresponding to the first discrimination phase (H_2_O-S1a). **E**) Paired data of the sniffing time of APP^WT^ and APP^Het^ mice corresponding to the second discrimination phase (S1c-S2a). **F**) Dispersion plots of APP^WT^ and APP^Het^ mice corresponding to the slope values of the first and second discrimination phases. **G)** Paired data of the sniffing time of young and aged wild type animals corresponding to the first habituation phase (S1a-S1b). **H**) Paired data of the sniffing time of the young and aged wild type animals corresponding to the second habituation phase (S2a-S2b). **I**) Dispersion plots of the slope values of young and aged mice corresponding to the first and second habituation phases. **J)** Paired data of the sniffing time of APP^WT^ and APP^Het^ mice corresponding to the first habituation phase (S1a-S1b). **K**) Paired data of the sniffing time of APP^WT^ and APP^Het^ mice corresponding to the second habituation phase (S2a-S2b). **L**) Dispersion plots of the slope values of APP^WT^ and APP^Het^ mice corresponding to the first and second habituation phases. Black dots in A, B, D, E, G, H, J, K and thick lines in C, F, I, L indicate mean ± SEM. Data were analyzed by one-way ANOVA with Tukey’s test. P ≤ 0.05 was considered statistically significant.

**Figure 7.**
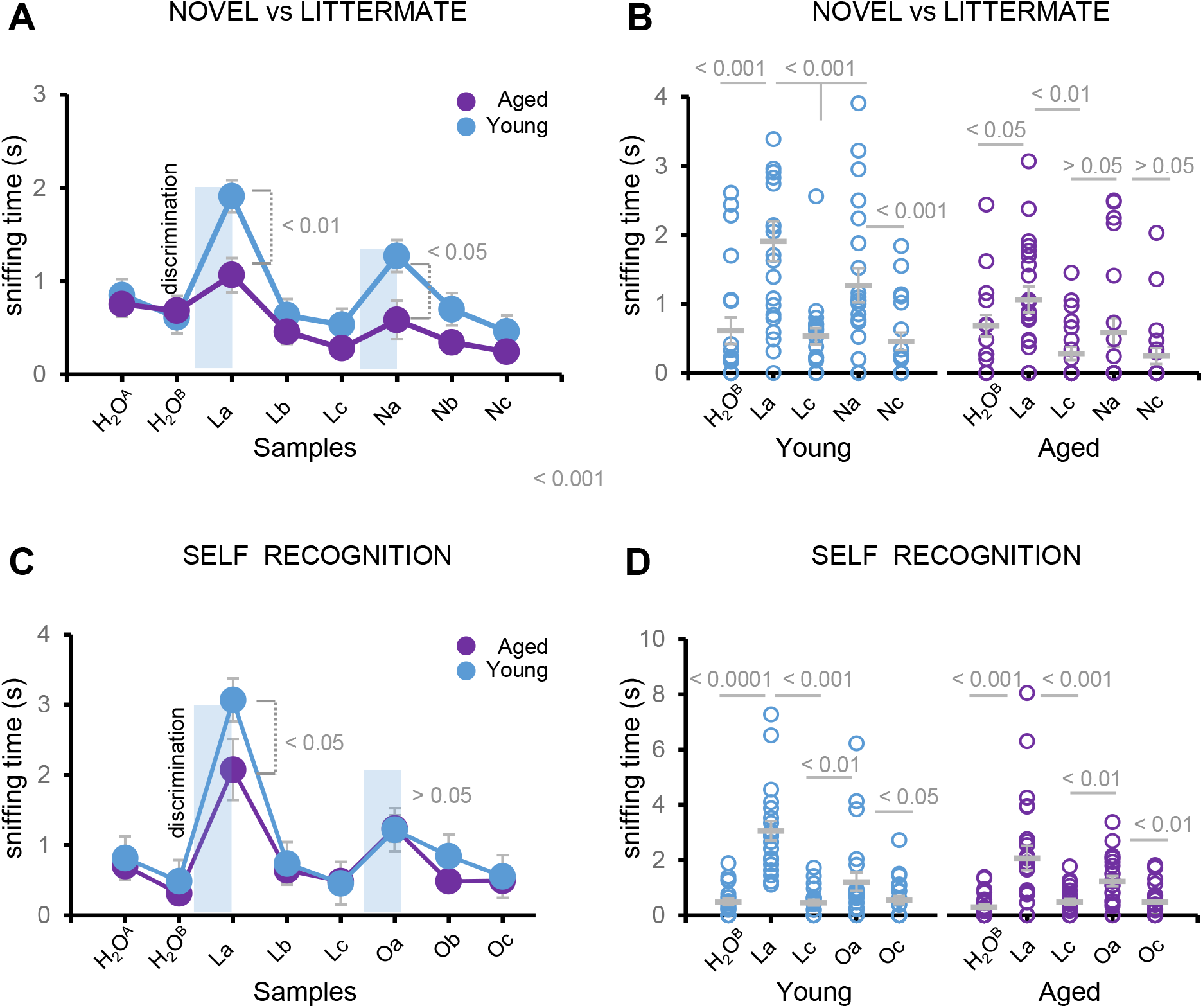
Age-related deficits in social discrimination and habituation are not influenced by previous experience. **A)** Average sniffing time of social habituation-dishabituation test in response to odors from novel (N) or littermate (L) subjects. **B)** Dispersion plot of the exploration time of aged and young animals in response to novel or littermate urine samples. **C)** Average sniffing time of social habituation-dishabituation test in response to littermate and the animal’s own urine (O). **D)** Dispersion plot of exploration time of aged and young animals in response to their own’s or littermate urine samples. Thick lines in dispersion plots indicate mean ± SEM. Number of mice: young: n = 21; aged: n = 23. Data were analyzed by a one-way ANOVA with Tukey’s test to test multiple comparisons with more than one variable. P ≤ 0.05 was considered statistically significant.

Since impairments in social odor discrimination were more significant in naturally aged mice, we asked whether these deficits also extended to the recognition of animal’s own odors, as the loss of self-awareness is a disrupting symptom of common occurrence in senescent subjects (La Joie et al., 2016, Valech et al., 2018). To this aim, animals were presented to samples of their own urine (O) during the second discrimination phase of the habituation-dishabituation test (**Fig. 7C, D**). Our data indicated that animal’s own urine was effective to elicit a typical discrimination (dishabituation) response, (**Fig. 7C, D**: sniffing time in seconds, **water control 1**: aged: 0.70 ± 0.10; young: 0.81 ± 0.10; **water control 2**: aged: 0.31 ± 0.09; young: 0.48 ± 0.11; **La**: aged: 2.07 ± 0.43; young: 3.06 ± 0.35; **Lb**: aged: 0.64 ± 0.16; young: 0.74 ± 0.17; **Lc**: aged: 0.48 ± 0.10; young: 0.45 ± 0.10; **Oa**: aged: 1.23 ± 0.18; young: 1.22 ± 0.33; **Ob**: aged: 0.48 ± 0.13; young: 0.84 ± 0.13; **Oc**: aged: 0.49 ± 0.11; young: 0.55 ± 0.15), suggesting that self-recognition is overall preserved in senescent mice.

### Social novelty is disrupted during pathological aging

Last, we explored whether the observed impairments in social odor exploration and discrimination may negatively impact social interaction in naturally aged and APP/PS1^Het^ mice. To this aim, we performed a three-chamber test (Nadler et al., 2004, Yang et al., 2011; **Fig. 8A**) to assess general sociability and social novelty in senescent and APP/PS1^Het^ animals. Our data indicated a reduction in social novelty in middle age APP1/PS1^Het^ mice, which was not detected in 2 year-old animals (**Fig. 8B-E, Sociability**, sniffing time in seconds, **E**: aged: 14.50 ± 1.90; young: 25.06 ± 2.96; APP/PS1^WT^: 39.15 ± 3.63; APP/PS1^Het^: 41.00 ± 4.07; **M1**^**A**^: aged: 47.24 ± 4.60; young: 64.35 ± 5.23; APP/PS1^WT^: 69.40 ± 6.05; APP/PS1^Het^: 69.73 ± 7.10; **Social novelty**, sniffing time in seconds, **M1**^**B**^: aged: 49.15 ± 4.20; young: 62.37 ± 5.01; APP/PS1^WT^: 71.30 ± 5.75; APP/PS1^Het^: 65.12 ± 6.70; **M2**: aged: 63.70 ± 5.30; young: 99.93 ± 9.83; APP/PS1^WT^: 93.07 ± 11.70; APP/PS1^Het^: 77.02 ± 8.43). Although senescent mice did not show significant impairments in either sociability or social novelty, they exhibited a mild increase in the latency to approach M1 during the sociability phase (**Additional file 7**), consistent with the overall decrease in the exploration time of social odors **(Additional files 2, 3)**. These findings exposed exacerbated deficits in an animal model of AD, suggesting a distinct impact of natural and pathological aging on social behavior.

**Figure 8.**
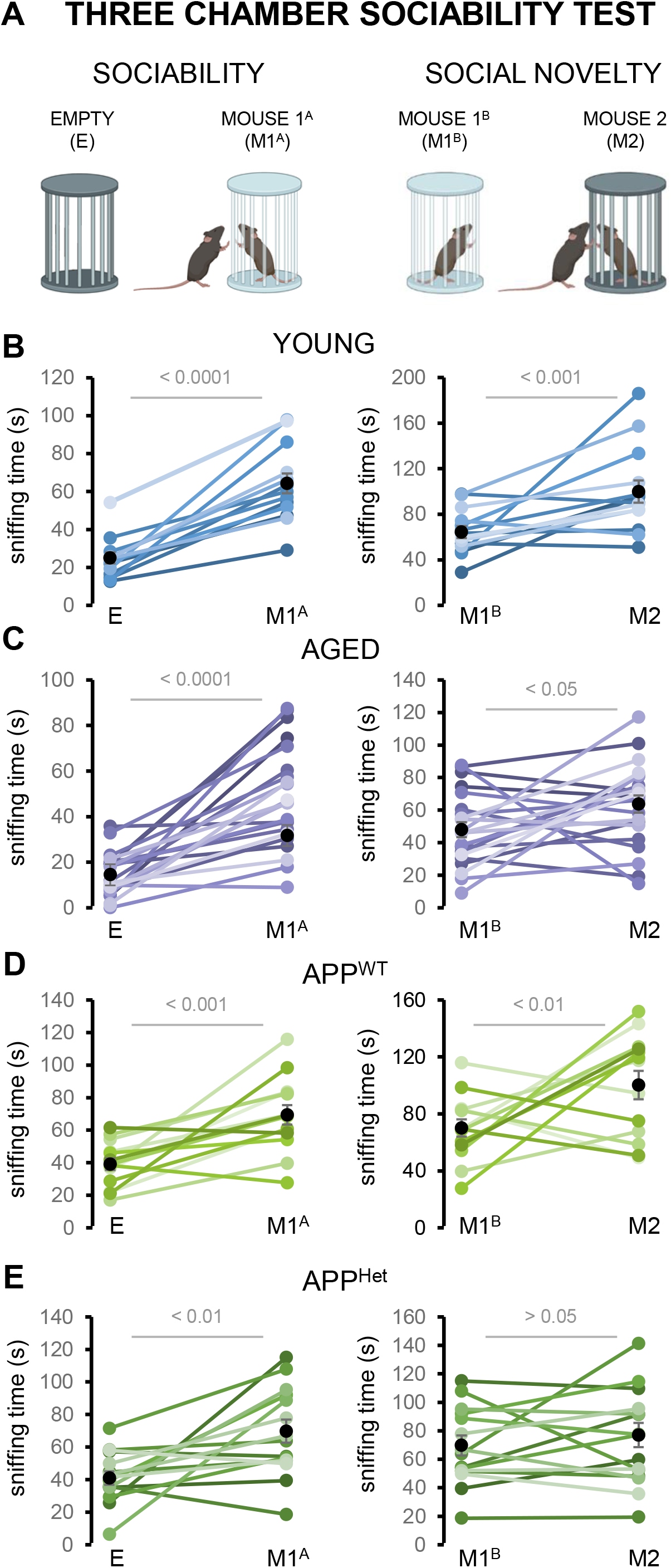
Social novelty is disrupted during pathological aging. **A)** Schematics of the three chamber test used in this study to test sociability and social novelty. In the sociability phase the sniffing time of E and M1^A^ are compared. Social novelty is estimated by quantifying the sniffing time of exploring M1^B^ versus M2 (see Methods). **B)** Paired data of the sniffing times of young wild type mice during the sociability (E-M1^A^) and social novelty (M1^B^-M2) phases. **C)** Paired data of the sniffing times of aged wild type mice during sociability and social novelty. **D)** Paired data of the sniffing times of middle age APP^WT^ mice during sociability) and social novelty. **E)** Paired data of the sniffing times of middle age APP^Het^ mice during sociability and social novelty. Black dots in the paired plots indicate mean ± SEM. Number of mice: young: n = 10; aged: n = 14; middle age control: n = 10; middle age APP/PS1^Het^: n = 12. Data were analyzed by a one-way ANOVA with Tukey’s test to test multiple comparisons with more than one variable. P ≤ 0.05 was considered statistically significant.

## DISCUSSION

The quality of social life has been proposed as a predictive factor for developing dementia or mental illness (Cohen et al., 1985, Stockdale et al., 2007, Wilson et al., 2007). However, the impairment of social functions with age is poorly understood with some authors suggesting that social deficits might be the consequence of generalized brain impairments (Johansson et al., 2018, Moran et al., 2012, Cavallini et al., 2013) or a symptom which might be developed independently of cognitive deficits (Spalletta et al., 2004, Mohs et al., 2000).

Our results revealed that both natural and pathological aging affect several key aspects of social information processing in mice including the exploration of social odors, social odor discrimination and habituation. To gain insight into the mechanisms underlying these deficits, we explored the age-related adaptions of the VSE, a central gateway for pheromone-encoded information in mammals (Jacobson, 1813, Halpern and Martínez-Marcos, 2003, Munger et al., 2009) and part of the accessory olfactory system whose aging process has been largely overlooked in contrast to other aspects of olfaction (Murphy, 2019).

Our data showed VSE alterations during natural aging but not in APP/PS1^Het^ mice, a standardized animal model of AD. Whereas the VSE of APP/PS1^Het^ mice maintained fairly stable neurogenic capabilities, senescent wild type animals showed a reduction in proliferative and stem cells (PCNA^+^ and Sox2^+^ cells) in the marginal VSE (**Fig. 2**) which in turn may explain the reduction of mature OMP^+^ neurons, sustentacular Sox2^+^ cells and the organ volume (**Fig. 1**). Our findings are consistent with two previous studies (Brann and Firestein, 2010, Mechin et al., 2021), which reported an overall thinning of the vomeronasal sensory epithelium (Mechin et al., 2021) and reduced VSE neurogenesis in the marginal zone of 2-year-old animals (Brann and Firestein, 2010). Here, we expand the current knowledge with novel information from a commonly used AD animal model (APP/PS1^Het^), revealing an unexpected preservation of the VSE proliferative capabilities in middle age APP/PS1^Het^ mice. These results contrast with the reduction of SVZ neurogenesis reported in similar animal models (Verret et al., 2007, Zhang et al., 2007, Zeng et al., 2016, Scopa et al., 2020), suggesting that VSE neurogenesis might be less susceptible to lesions (Brann and Firestein, 2010) and pathological conditions than other proliferative areas. However, we cannot rule out the possibility that the VSE of APP/PS1^Het^ animals may develop potential alterations at a later age (> 1 year-old). An aspect that we were prevented to investigate due to the short life expectancy (∼ 14-16 months-old) of this mouse model.

A relevant aspect of our work is that most previous studies addressing olfactory decline employed synthetic or neutral odors, thus remaining uncertain the effect of healthy and diseased aging in the recognition of social cues. To gain insight into this question, we investigated the exploration time and the habituation-dishabituation response to social odors (urine) (**Figs. 3, 5, 6**). Our findings revealed that despite the distinctive effects of natural and pathological aging on the VSE, both processes impaired the exploration of socio-sexual cues, social discrimination-habituation, and social behavior, suggesting fundamental differences in the mechanisms by which healthy and diseased aging impact social information processing.

Further experiments are needed to establish causality, but our results suggest that the observed VSE alterations in senescent mice underlie deficits in urine exploration time (**Fig. 3, Additional files 2, 3**) which could impair the processing of social information. VSE volume data from middle age APP/PS1^WT^ control animals suggest that VSE structural changes might appear around 1 year-old of age (**Fig. 1A**: VES volume mm^3^: 4 month-old: 0.190 ± 0.008; 1-year-old APP/PS1^WT^: 0.170 ± 0.007; p = 0.03; two-way ANOVA with Tuckey’s test), although their functional consequences might not become apparent until advanced stages of aging (2-year-old). This scenario suggests a parsimonious VNO decay that matches the aging of other olfactory areas like the OB. Several studies have shown that despite the reduced regeneration rate of olfactory sensory neurons (OSNs), OB alterations involving a reduced number of synaptic contacts (Richard et al., 2010), expression of odorant receptor genes (Lee et al., 2009; Khan et al., 2013) and changes in the OSNs dynamic range (Kass et al., 2018) just become apparent in 2-year-old mice. This evidence suggests that although age-related changes may start to appear earlier (Mobley et al., 2014), functional deficits might exhibit a late onset, suggesting compensatory mechanisms to preserve the processing of olfactory cues relevant for the survival of aged animals (Tobiansky et al., 2012, Tikhonova et al., 2015). In contrast to natural steady decline, pathological conditions may accelerate functional deficits even in the absence of detectable modifications of peripheral organs, suggesting impairments in the central processing of social information. As such, defects in the exploration of social odors were found exacerbated in middle age APP/PS1^Het^ mice (**Fig. 3D, E**), a condition likely to aggravate frailty and reduce life expectancy in these animals (as observed for APP/PSEN1^Het^ in our experimental conditions).

Furthermore, a detailed analysis of the first discrimination phase during the modified social habituation-dishabituation test (presentation of S1a urine sample after 24 hours; **Fig. 5F, G**) revealed significant impairments in both 2-year old wild type animals and middle age APP/PS1^Het^ mice, suggesting that in parallel to peripheral sensory decline, the downstream pathways involved in social cue recognition may be affected during both natural and pathological aging. To control for a potential contribution of novelty in the social habituation-dishabituation test, we presented urine samples from littermate and strange animals finding similar deficits in discrimination (**Fig. 7**). Interestingly, although senescent animals showed severe discrimination deficits they preserved the ability to differentiate between urine from a novel subject and their own, preventing to explore further the underlying mechanisms of subjective perception loss, a disrupting symptom of senescence and dementia which currently lacks appropriate animal models for preclinical studies (Jessen et al., 2020).

Age-related decline of olfactory detection is likely to occur due to alterations of both the main olfactory epithelium and the VNO (Apfelbach et al., 1991, Lee et al., 2009, Ueha et al., 2018), thus potentially affecting other odor modalities. Here, we sought to explore this question by performing odor-evoked sniffing tests using social and neutral synthetic odors (**Fig. 4**). Quantification of the exploration time across various IA dilutions exposed equivalent measurements in senescent and young mice, and even enhanced responses in APP/PS1^Het^ mice. Furthermore, results from a FFT, revealed no significant differences in the latency time to find the hidden food pellets (**Fig. 4E**) indicating that defects in social odor detection related to natural and pathological aging may be more severe than other odor modalities. A plausible explanation for the specific impairment in the exploration of social odors may involve the reduction of reproductive activity in old animals, but not in foraging and feeding behaviors (Harb et al., 2014). Furthermore, the decline to detect specific odors may rise from alterations on the number of sensory neurons and synaptic connections specifically related to the discrimination of conspecifics (Bielsky et al., 2005, Wacker et al., 2010, Tobin et al., 2010) and heterospecific social cues (e.g. predator) (Kang et al., 2009, Wacker et al., 2010). In particular, impairments in social odor exploration and discrimination may rise from maladaptations of the VNO-AOB axis, consistent with the observed decrease in AOB volume (**Fig. 1B**). Future studies should deepen into the anatomical and functional properties of the VNO-AOB circuit in order to obtain a complete picture on how healthy and pathological aging affect social information processing.

Finally, we sought to determine how the age-related defects in social odor sensitivity, discrimination and memory shaped social behavior. Results from a three chamber test (**Fig. 8**) indicated that sociability was overall preserved in senescent and APP/PS1^Het^ animals. In contrast, social novelty was severely impaired in APP/PS1^Het^ mice consistently with previous studies (Cheng et al., 2014, Olesen et al., 2016, Shoji and Miyakawa, 2019, Locci et al., 2021). Deficits in social novelty in APP/PS1^Het^ mice could be linked to the deficits in social discrimination observed in the habituation-dishabituation test (**Fig. 5D, E** and **Fig. 6D-F**) which could impair the recognition of M2 as a novel subject (Enwere et al., 2004, Patel and Larson, 2009, Moreno et al., 2014). Consistent with the overall decrease in the exploration time of social odors **(Fig. 3, Additional files 2, 3)**, aged animals exhibited a mild increase in the latency to approach the novel mouse (M1) during the first sociability phase (**Additional file 7**). These findings indicate that despite the reduction in social odor exploration and discrimination, overall sociability and novelty are majorly preserved in naturally aged mice. In contrast, the APP/PS1 neurodegenerative model shows a measurable impairment in social preference. This evidence opens the intriguing possibility that neuronal circuits underlying specific social functions (sociability vs social memory) may be particularly susceptible to pathological aging.

## CONCLUSION

Olfactory deficits are a common symptom of natural and pathological aging. While multiple aged-related changes of the olfactory sensory epithelium are described, the aging of the pheromone detection system, a major gateway for social information, has been largely overlooked. This study reveals that whereas natural aging elicits a decline in VSE cell proliferation, likely to reduce the number of mature sensory neurons and the overall volume of the organ, a common animal model of AD exhibits normal proliferation capacities and no obvious morphological alterations. Despite their distinctive effects at the cellular level, both natural and pathological aging disrupt the detection of social odors in a more severe way than other odor modalities. Furthermore, social detection and social behavior impairments were exacerbated in APP/PS1^Het^ mice, indicating pathological aging may impact the downstream processing of social information even in the absence of obvious alterations of the peripheral organs.

## DECLARATIONS

### Availability of data and materials

The datasets used and/or analysed during the current study are available from the corresponding author on reasonable request.

### Author contributions

The project was led by S.J. Brain sample preparation, data acquisition, and analysis was done by A.P. Result interpretation, manuscript preparation, and editing was done by S.J., P.C. and A.P. with help of members from the Jurado Lab at the Instituto de Neurociencias (CSIC-UMH).

## Acknowledgements

We are grateful to the teams of the Imaging and Animal Facilities at the Institute of Neuroscience, and all members of the Jurado and Chamero laboratories for their support during the realization of this work.

## FIGURE LEGENDS

**Additional file 1. Structural modifications of the mouse VSE and AOB during natural and pathological aging. A)** Schematics depicting the VNO divisions into marginal (M), intermediate (I) and central (C). **B**) Dispersion plots of the number of OMP^+^ cells in the marginal zone of the VSE combining anterior, medial and posterior sections. Thick lines indicate the mean ± SEM. Data were obtained from 10-16 slices from at least three animals per condition. Statistical comparisons were calculated by two-way ANOVA with Tukey’s test. P ≤ 0.05 was considered statistically significant.

**Additional file 2. Social odor exploration time during natural aging**. Dispersion plots represent the raw sniffing time of 2, 6, 12 and 24 months-old animals for the various serial dilutions tested in **Fig. 3**. Thick grey lines indicate mean ± SEM. Data were analyzed by a one-way ANOVA with Tukey’s test to test multiple comparisons with more than one variable. P and n values are indicated in the tables. P ≤ 0.05 was considered statistically significant.

**Additional file 3. Statistical comparison between different urine dilutions for each age tested (2, 6, 12 and 24 months-old)**. Dispersion plots represent the sniffing time and trend curves of 2, 6, 12 and 24 months-old animals. Thick dots indicate mean ± SEM. P values for each statistical comparison are indicated in the tables on the right. Data were analyzed by a one-way ANOVA with Tukey’s test to test multiple comparisons with more than one variable. P ≤ 0.05 was considered statistically significant.

**Additional files 4. Aged-related deficits in social odor exploration exhibit no significant sex differences. A**) Average sniffing time of urine serial dilutions of adult and aged female wild type mice. **B**) Average exploration (sniffing) time of urine serial dilutions of adult and aged male wild type mice. **C**) Dispersion plots of the sniffing time of urine dilutions that show significant impairments in female mice (nd, non-diluted; 1:10; 1:500; 1:1000). **D**) Dispersion plots of the sniffing time of the urine dilutions that show significant changes in male mice (nd, non-diluted; 1:250; 1:500; 1:1000). Plots indicate mean ± SEM. Number of mice: adult females: n = 13; adult males: n = 11; aged females: n = 14; aged males: n = 11. Data were analyzed by a one-way ANOVA with Tukey’s test to test multiple comparisons with more than one variable. P ≤ 0.05 was considered statistically significant.

**Additional file 5. Social odor exploration of APP**^**WT**^ **and APP**^**Het**^ **mice**. Dispersion plots represent the raw sniffing time of middle age APP^WT^ and APP^Het^ mice for the various serial dilutions. Thick grey lines indicate mean ± SEM. Number of mice for each condition is indicated in the table below. Data were analyzed by a one-way ANOVA with Tukey’s test to test multiple comparisons with more than one variable. P ≤ 0.05 was considered statistically significant.

**Additional file 6. Statistical comparison between different urine dilutions of APP/PS1**^**WT**^ **and APP/PS1**^**Het**^ **mice**. Dispersion plots represent the sniffing time and trend curves of middle age APP^WT^ and APP^Het^ mice for the various serial dilutions. Thick dots indicate mean ± SEM. P values for each statistical comparison are indicated in the tables on the right. Data were analyzed by a one-way ANOVA with Tukey’s test to test multiple comparisons with more than one variable. P ≤ 0.05 was considered statistically significant.

**Additional file 7. Latency to approach the first stranger (M1**^**A**^**) in the three chamber sociability test is increased in naturally aged animals. A**) The sociability phase of the three chamber test involves exposing the test animal to a stranger mouse (M1^A^) for the first time, which elicits an increase in the exploration time in that side of the chamber. **B**) Dispersion plots of the latency to approach M1 during the first encounter of the sociability phase. Thick lines indicate mean ± SEM. Data were analyzed by one way-ANOVA with Tukey’s test. P ≤ 0.05 was considered statistically significant.

## ABBREVIATIONS

AD: Alzheimer’s disease
AOB: Anterior olfactory bulb
APP: Amyloid precursor protein
C: VNO central zone
CTF: Corrected total fluorescence
FFT: Food finding test
I: VNO intermediate zone
IA: Isoamyl acetate
M: VNO marginal zone
MOB: Main olfactory bulb
MOE: Main olfactory epithelium
NSE: Non-sensory epithelium
OCT: Optimal cutting temperature
OMP: Olfactory marker protein
PCNA: Proliferative cell nuclear antigen
PS1: Presinilin-1
SCL: Supporting cell layer
Sox2: SRY-box transcription factor 2
VL: Vomeronasal lumen
VNO: Vomeronasal organ
VSE: Vomeronasal epithelium

