## Additional file for "Natural and pathological aging distinctively impact the vomeronasal detection system and social behavior"

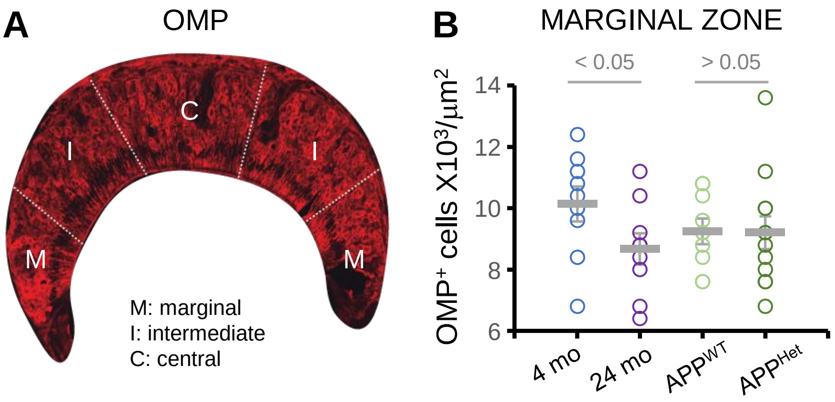


Additional file 1

### SOCIAL ODOR EXPLORATION – NATURAL AGING

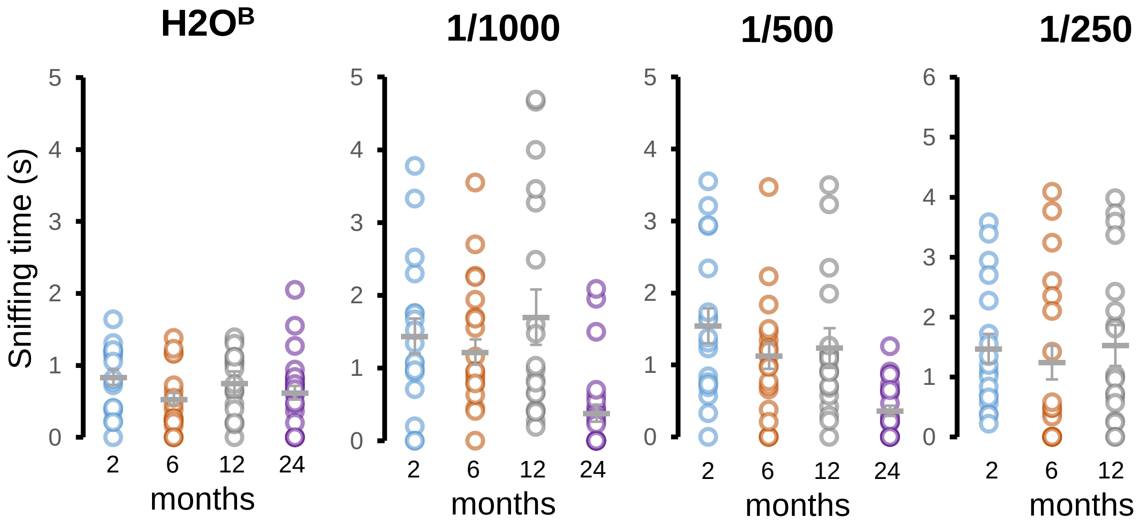


| H2O^B | |
|---|---|
| **Comparison** | ***P*** |
| 2 m vs 6 m | **0.02** |
| 2 m vs 12 m | ns |
| 2 m vs 24 m | ns |
| 6 m vs 12 m | ns |
| 6 m vs 24 m | ns |
| 12 m vs 24 m | ns |

| 1:1000 urine dilution | |
|---|---|
| **Comparison** | ***P*** |
| 2 m vs 6 m | ns |
| 2 m vs 12 m | ns |
| 2 m vs 24 m | **0.0007** |
| 6 m vs 12 m | ns |
| 6 m vs 24 m | **0.0004** |
| 12 m vs 24 m | **0.002** |

| 1:500 urine dilution | |
|---|---|
| **Comparison** | ***P*** |
| 2 m vs 6 m | ns |
| 2 m vs 12 m | ns |
| 2 m vs 24 m | **0.0002** |
| 6 m vs 12 m | ns |
| 6 m vs 24 m | **0.0005** |
| 12 m vs 24 m | **0.0008** |

| 1:250 urine dilution | |
|---|---|
| **Comparison** | ***P*** |
| 2 m vs 6 m | ns |
| 2 m vs 12 m | ns |
| 2 m vs 24 m | **0.03** |
| 6 m vs 12 m | ns |
| 6 m vs 24 m | ns |
| 12 m vs 24 m | **0.04** |

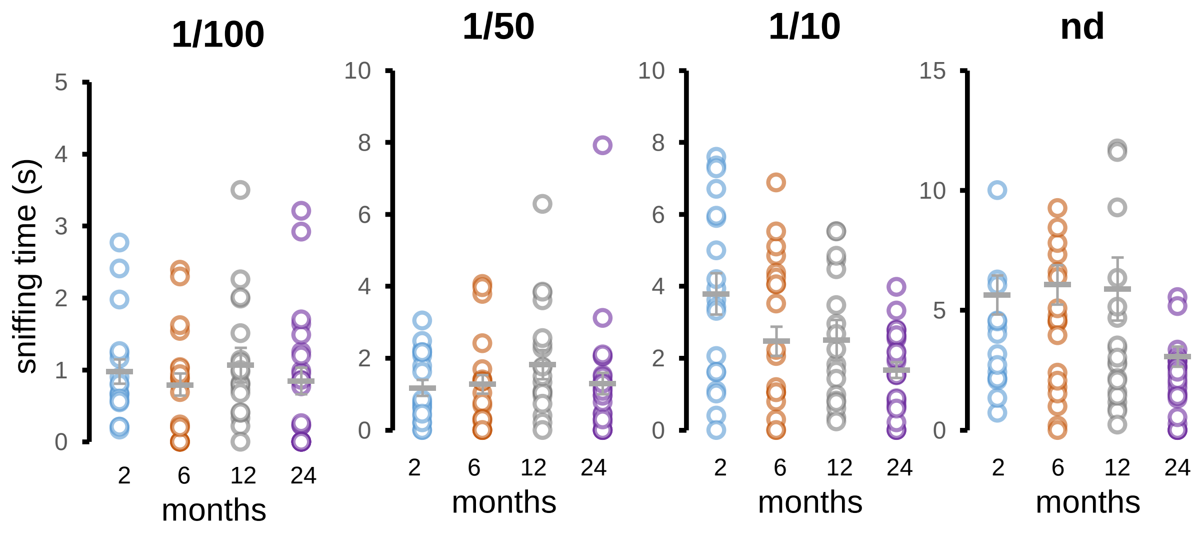


| 1:100 urine dilution | |
|---|---|
| **Comparison** | ***P*** |
| 2 m vs 6 m | ns |
| 2 m vs 12 m | ns |
| 2 m vs 24 m | ns |
| 6 m vs 12 m | ns |
| 6 m vs 24 m | ns |
| 12 m vs 24 m | ns |

| 1:50 urine dilution | |
|---|---|
| **Comparison** | ***P*** |
| 2 m vs 6 m | ns |
| 2 m vs 12 m | ns |
| 2 m vs 24 m | ns |
| 6 m vs 12 m | ns |
| 6 m vs 24 m | ns |
| 12 m vs 24 m | ns |

| 1:10 urine dilution | |
|---|---|
| **Comparison** | ***P*** |
| 2 m vs 6 m | **0.08** |
| 2 m vs 12 m | **0.04** |
| 2 m vs 24 m | **0.002** |
| 6 m vs 12 m | ns |
| 6 m vs 24 m | ns |
| 12 m vs 24 m | ns |

| Non diluted | |
|---|---|
| **Comparison** | ***P*** |
| 2 m vs 6 m | ns |
| 2 m vs 12 m | ns |
| 2 m vs 24 m | **0.04** |
| 6 m vs 12 m | ns |
| 6 m vs 24 m | **0.003** |
| 12 m vs 24 m | **0.02** |

| dilution / age | ***$H_2O^B$*** | ***1:000*** | ***1:500*** | ***1:250*** | ***1.100*** | ***1:50*** | ***1:10*** | ***nd*** |
|---|---|---|---|---|---|---|---|---|
| | ***n*** | **n** | ***n*** | ***n*** | ***n*** | ***n*** | ***n*** | ***n*** |
| 2 months | 18 | 18 | 18 | 18 | 18 | 19 | 19 | 15 |
| 6 months | 24 | 19 | 20 | 20 | 19 | 21 | 20 | 21 |
| 12 months | 20 | 19 | 20 | 19 | 20 | 20 | 18 | 19 |
| 24 months | 26 | 25 | 25 | 25 | 26 | 27 | 26 | 26 |

Additional file 2

### SOCIAL ODOR EXPLORATION – NATURAL AGING

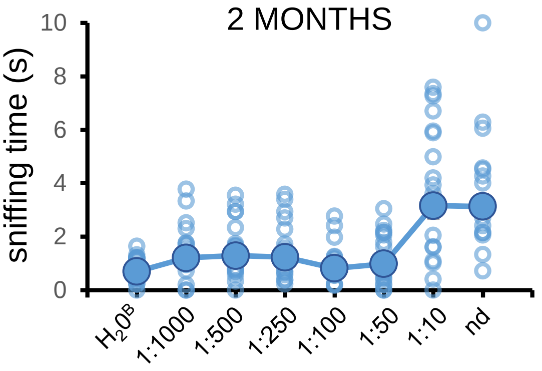


**One Way ANOVA – Tukey test**

| 2 months | |
|---|---|
| **Comparison** | ***P*** |
| $H_2O^B$ vs 1/1000 | 0.91 |
| $H_2O^B$ vs 1/500 | 0.81 |
| 1/1000 vs 1/500 | 1.00 |
| $H_2O^B$ vs 1/250 | 0.88 |
| 1/250 vs 1/1000 | 1.00 |
| 1/250 vs 1/500 | 1.00 |
| $H_2O^B$ vs 1/100 | 0.99 |
| 1/100 vs 1/1000 | 0.97 |
| 1/100 vs 1/500 | 0.93 |
| 1/100 vs 1/250 | 0.99 |
| $H_2O^B$ vs 1/50 | 0.99 |
| 1/50 vs 1/1000 | 0.99 |
| 1/500 vs 1/50 | 0.99 |
| 1/250 vs 1/50 | 0.99 |

| 2 months | |
|---|---|
| **Comparison** | ***P*** |
| 1/100 vs 1/50 | 0.99 |
| $H_2O^B$ vs 1/10 | **$1.5 x10^{-7}$** |
| 1/10 vs 1/1000 | **$5.0 x10^{-5}$** |
| 1/10 vs 1/500 | **$1.3 x10^{-4}$** |
| 1/10 vs 1/250 | **$7.0 x10^{-5}$** |
| 1/10 vs 1/100 | **$5.8 x10^{-7}$** |
| 1/10 vs 1/50 | **$3.0 x10^{-6}$** |
| Nd vs $H_2O^B$ | **$1.0 x10^{-6}$** |
| Nd vs 1/1000 | **$2.2 x10^{-4}$** |
| Nd vs 1/500 | **$5.2 x10^{-4}$** |
| Nd vs 1/250 | **$3.0 x10^{-4}$** |
| Nd vs 1/100 | **$4.0 x10^{-6}$** |
| Nd vs 1/50 | **$1.8 x10^{-5}$** |
| Nd vs 1/10 | 1.00 |

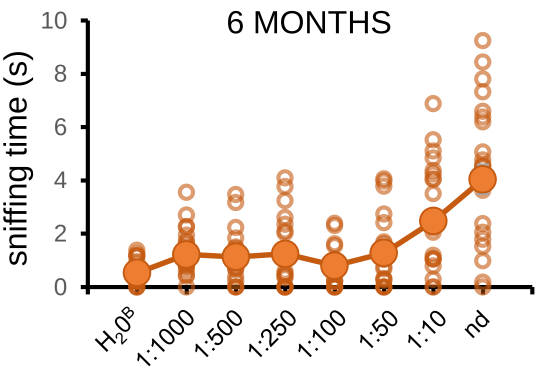


**One Way ANOVA – Tukey test**

| 6 months | |
|---|---|
| **Comparison** | ***P*** |
| $H_2O^B$ vs 1/1000 | 0.63 |
| $H_2O^B$ vs 1/500 | 0.85 |
| 1/1000 vs 1/500 | 0.99 |
| $H_2O^B$ vs 1/250 | 0.82 |
| 1/250 vs 1/1000 | 0.99 |
| 1/250 vs 1/500 | 1.00 |
| $H_2O^B$ vs 1/100 | 0.99 |
| 1/100 vs 1/1000 | 0.94 |
| 1/100 vs 1/500 | 0.99 |
| 1/100 vs 1/250 | 0.98 |
| $H_2O^B$ vs 1/50 | 0.53 |
| 1/50 vs 1/1000 | 1.0 |
| 1/500 vs 1/50 | 0.99 |
| 1/250 vs 1/50 | 0.99 |

| 6 months | |
|---|---|
| **Comparison** | ***P*** |
| 1/100 vs 1/50 | 0.90 |
| $H_2O^B$ vs 1/10 | **$6.4 x10^{-5}$** |
| 1/10 vs 1/1000 | 0.07 |
| 1/10 vs 1/500 | **0.02** |
| 1/10 vs 1/250 | **0.03** |
| 1/10 vs 1/100 | **0.0016** |
| 1/10 vs 1/50 | 0.07 |
| Nd vs $H_2O^B$ | **$2.2 x10^{-8}$** |
| Nd vs 1/1000 | **$8.3 x10^{-8}$** |
| Nd vs 1/500 | **$4.1 x10^{-8}$** |
| Nd vs 1/250 | **$4,3 x10^{-8}$** |
| Nd vs 1/100 | **$4.5 x10^{-8}$** |
| Nd vs 1/50 | **$7.0 x10^{-8}$** |
| Nd vs 1/10 | **0.01** |

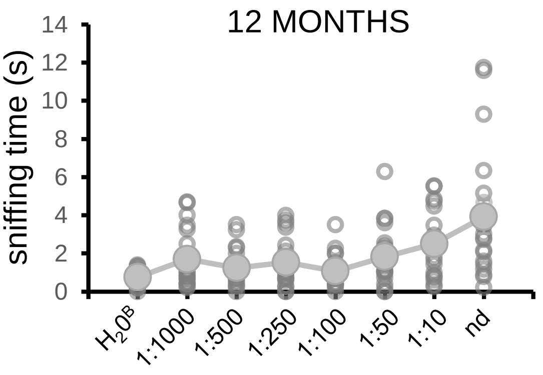


**One Way ANOVA – Tukey test**

| 12 months | |
|---|---|
| **Comparison** | ***P*** |
| $H_2O^B$ vs 1/1000 | 0.67 |
| $H_2O^B$ vs 1/500 | 0.98 |
| 1/1000 vs 1/500 | 0.99 |
| $H_2O^B$ vs 1/250 | 0.85 |
| 1/250 vs 1/1000 | 0.99 |
| 1/250 vs 1/500 | 0.99 |
| $H_2O^B$ vs 1/100 | 0.99 |
| 1/100 vs 1/1000 | 0.94 |
| 1/100 vs 1/500 | 0.99 |
| 1/100 vs 1/250 | 0.99 |
| $H_2O^B$ vs 1/50 | 0.50 |
| 1/50 vs 1/1000 | 1.0 |
| 1/500 vs 1/50 | 0.95 |
| 1/250 vs 1/50 | 0.99 |

| 12 months | |
|---|---|
| **Comparison** | ***P*** |
| 1/100 vs 1/50 | 0.85 |
| $H_2O^B$ vs 1/10 | 0.04 |
| 1/10 vs 1/1000 | 0.84 |
| 1/10 vs 1/500 | 0.31 |
| 1/10 vs 1/250 | 0.67 |
| 1/10 vs 1/100 | 0.17 |
| 1/10 vs 1/50 | 0.92 |
| Nd vs $H_2O^B$ | **$1.3 x10^{-6}$** |
| Nd vs 1/1000 | 0.002 |
| Nd vs 1/500 | **$7.2 x10^{-5}$** |
| Nd vs 1/250 | **$8.0 x10^{-4}$** |
| Nd vs 1/100 | **$2.0 x10^{-5}$** |
| Nd vs 1/50 | 0.005 |
| Nd vs 1/10 | 0.20 |

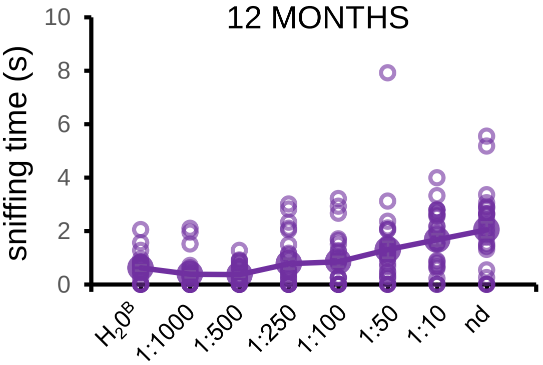


**One Way ANOVA – Tukey test**

| 24 months | |
|---|---|
| **Comparison** | ***P*** |
| $H_2O^B$ vs 1/1000 | 0.99 |
| $H_2O^B$ vs 1/500 | 0.98 |
| 1/1000 vs 1/500 | 1.0 |
| $H_2O^B$ vs 1/250 | 0.99 |
| 1/250 vs 1/1000 | 0.87 |
| 1/250 vs 1/500 | 0.84 |
| $H_2O^B$ vs 1/100 | 0.99 |
| 1/100 vs 1/1000 | 0.73 |
| 1/100 vs 1/500 | 0.69 |
| 1/100 vs 1/250 | 1.0 |
| $H_2O^B$ vs 1/50 | 0.25 |
| 1/50 vs 1/1000 | 0.03 |
| 1/500 vs 1/50 | 0.03 |
| 1/250 vs 1/50 | 0.6 |

| 24 months | |
|---|---|
| **Comparison** | ***P*** |
| 1/100 vs 1/50 | 0.76 |
| $H_2O^B$ vs 1/10 | 0.007 |
| 1/10 vs 1/1000 | **$3.4 x10^{-4}$** |
| 1/10 vs 1/500 | **$2.6 x10^{-4}$** |
| 1/10 vs 1/250 | 0.04 |
| 1/10 vs 1/100 | 0.08 |
| 1/10 vs 1/50 | 0.88 |
| Nd vs $H_2O^B$ | **$3.4 x10^{-5}$** |
| Nd vs 1/1000 | **$8.4 x10^{-7}$** |
| Nd vs 1/500 | **$6.3 x10^{-7}$** |
| Nd vs 1/250 | **$4,6 x10^{-4}$** |
| Nd vs 1/100 | 0.001 |
| Nd vs 1/50 | 0.14 |
| Nd vs 1/10 | 0.89 |

### SOCIAL ODOR EXPLORATION

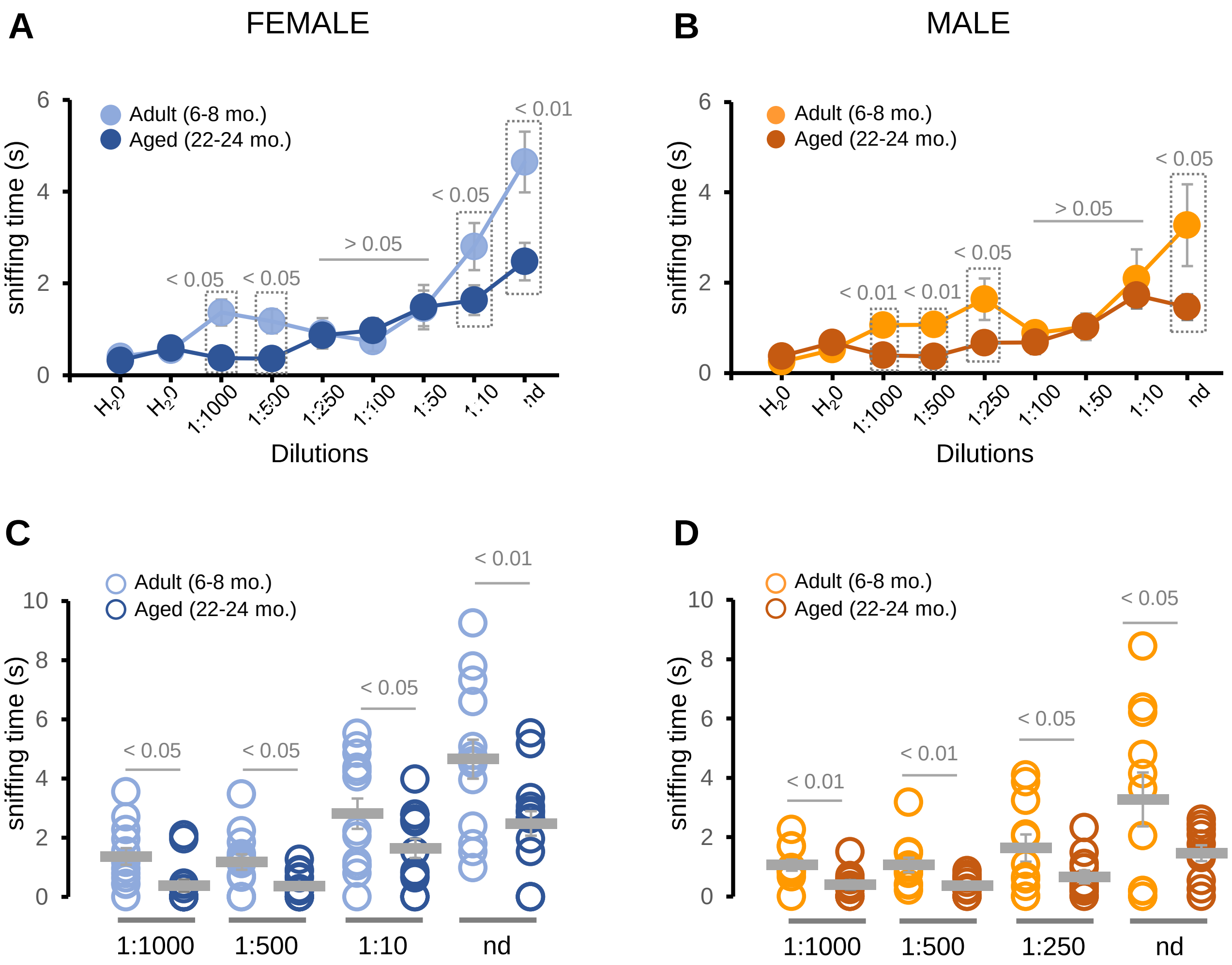

### SOCIAL ODOR EXPLORATION - PATHOLOGICAL AGING

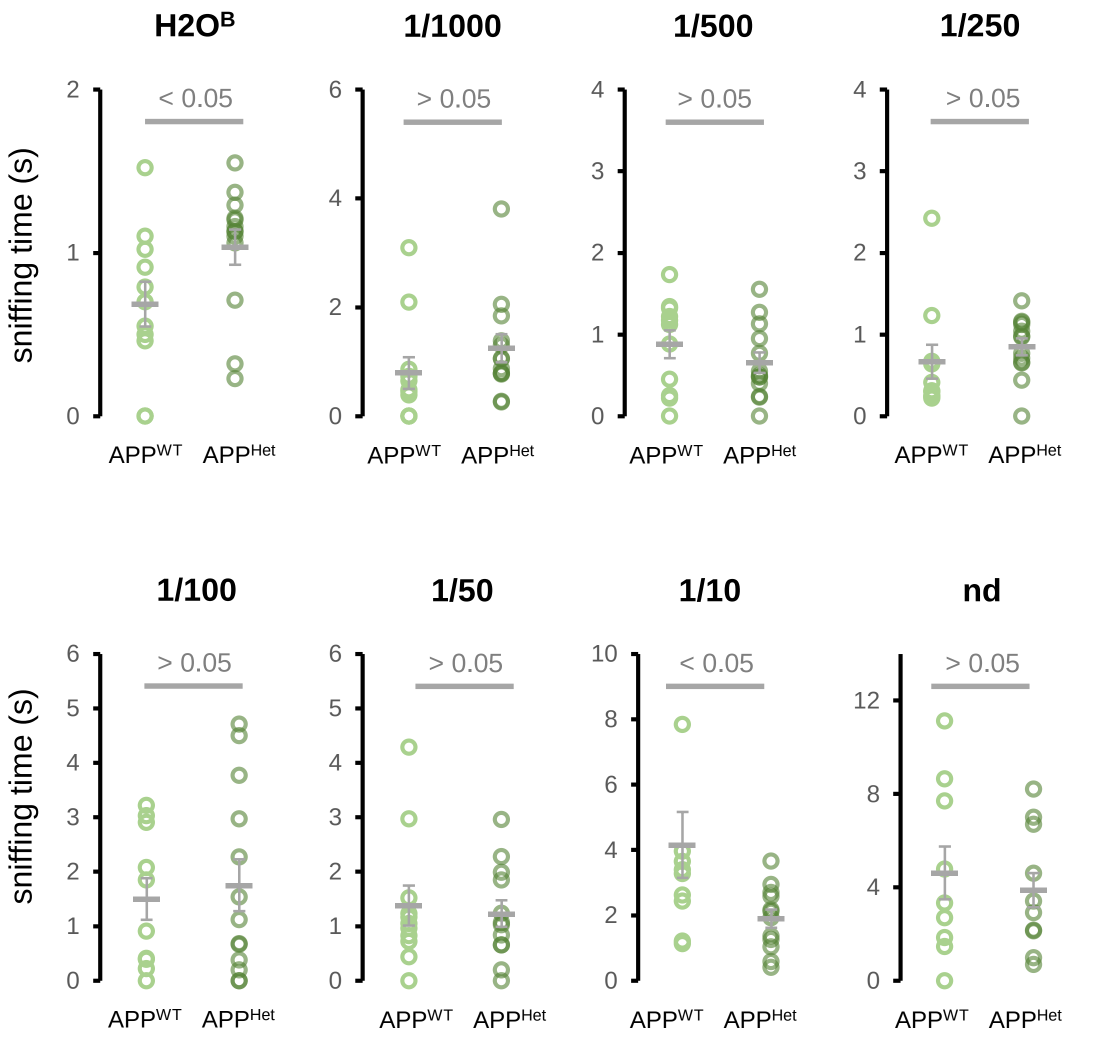


| dilution / genotype | $H_2O_B$ | 1:1000 | 1:500 | 1:250 | 1.100 | 1:50 | 1:10 | nd |
|---|---|---|---|---|---|---|---|---|
| | n | n | n | n | n | n | n | n |
| APP[WT] | 11 | 11 | 11 | 10 | 10 | 11 | 10 | 9 |
| APP[Het] | 13 | 13 | 13 | 13 | 13 | 12 | 12 | 10 |

### SOCIAL ODOR EXPLORATION – PATHOLOGICAL AGING

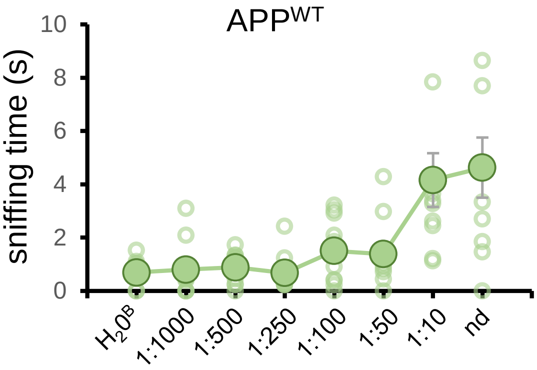


**One Way ANOVA – Tukey test**

| **APP$^{WT}$** | |
|---|---|
| **Comparison** | ***P*** |
| $H_2O^B$ vs 1/1000 | 1.00 |
| $H_2O^B$ vs 1/500 | 1.00 |
| 1/1000 vs 1/500 | 1.00 |
| $H_2O^B$ vs 1/250 | 1.00 |
| 1/250 vs 1/1000 | 1.00 |
| 1/250 vs 1/500 | 0.99 |
| $H_2O^B$ vs 1/100 | 0.97 |
| 1/100 vs 1/1000 | 0.99 |
| 1/100 vs 1/500 | 0.99 |
| 1/100 vs 1/250 | 0.97 |
| $H_2O^B$ vs 1/50 | 0.98 |
| 1/50 vs 1/1000 | 0.99 |
| 1/500 vs 1/50 | 0.99 |
| 1/250 vs 1/50 | 0.98 |

| **APP$^{WT}$** | |
|---|---|
| **Comparison** | ***P*** |
| 1/100 vs 1/50 | 1.00 |
| $H_2O^B$ vs 1/10 | 0.001 |
| 1/10 vs 1/1000 | 0.002 |
| 1/10 vs 1/500 | 0.003 |
| 1/10 vs 1/250 | 0.002 |
| 1/10 vs 1/100 | 0.04 |
| 1/10 vs 1/50 | 0.02 |
| Nd vs $H_2O^B$ | **3.1 x10$^{-4}$** |
| Nd vs 1/1000 | **5.0 x10$^{-4}$** |
| Nd vs 1/500 | **7.2 x10$^{-4}$** |
| Nd vs 1/250 | **4.1 x10$^{-4}$** |
| Nd vs 1/100 | 0.01 |
| Nd vs 1/50 | 0.005 |
| Nd vs 1/10 | 0.99 |

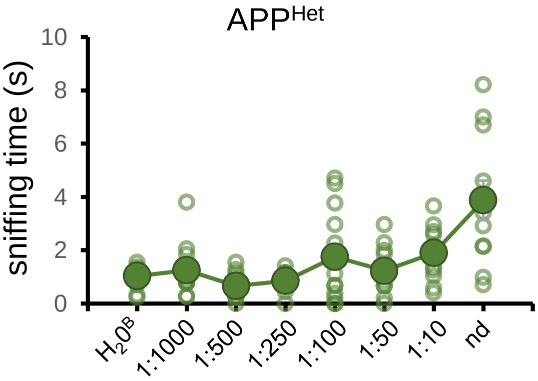


**One Way ANOVA – Tukey test**

| **APP$^{Het}$** | |
|---|---|
| **Comparison** | ***P*** |
| $H_2O^B$ vs 1/1000 | 0.99 |
| $H_2O^B$ vs 1/500 | 0.99 |
| 1/1000 vs 1/500 | 0.91 |
| $H_2O^B$ vs 1/250 | 0.99 |
| 1/250 vs 1/1000 | 0.99 |
| 1/250 vs 1/500 | 0.99 |
| $H_2O^B$ vs 1/100 | 0.79 |
| 1/100 vs 1/1000 | 0.96 |
| 1/100 vs 1/500 | 0.30 |
| 1/100 vs 1/250 | 0.99 |
| $H_2O^B$ vs 1/50 | 0.99 |
| 1/50 vs 1/1000 | 1.00 |
| 1/500 vs 1/50 | 0.93 |
| 1/250 vs 1/50 | 0.99 |

| **APP$^{Het}$** | |
|---|---|
| **Comparison** | ***P*** |
| 1/100 vs 1/50 | 0.95 |
| $H_2O^B$ vs 1/10 | 0.63 |
| 1/10 vs 1/1000 | 0.88 |
| 1/10 vs 1/500 | 0.18 |
| 1/10 vs 1/250 | 0.38 |
| 1/10 vs 1/100 | 0.99 |
| 1/10 vs 1/50 | 0.90 |
| Nd vs $H_2O^B$ | **6.6 x10$^{-6}$** |
| Nd vs 1/1000 | **3.7 x10$^{-5}$** |
| Nd vs 1/500 | **3.4 x10$^{-7}$** |
| Nd vs 1/250 | **1.5 x10$^{-6}$** |
| Nd vs 1/100 | 0.002 |
| Nd vs 1/50 | **4.7 x10$^{-5}$** |
| Nd vs 1/10 | 0.005 |

### THREE CHAMBER SOCIABILITY TEST

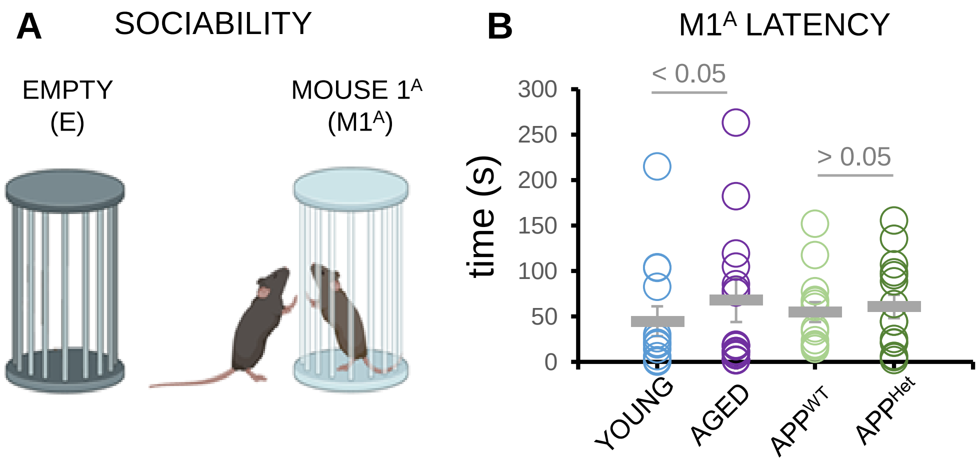


Additional file 7
